# Origin of the type I antifreeze gene in flounders in response to Cenozoic climate change

**DOI:** 10.1101/2021.09.21.461085

**Authors:** Laurie A. Graham, Sherry Y. Gauthier, Peter L. Davies

## Abstract

Antifreeze proteins (AFPs) inhibit ice growth within fish and protect them from freezing in icy seawater. Alanine-rich, alpha-helical AFPs (type I) have independently (convergently) evolved in four branches of fishes, one of which is a subsection of the righteye flounders. The origin of this gene family has been elucidated by sequencing two loci from a starry flounder, *Platichthys stellatus*, collected off Vancouver Island, British Columbia. The first locus had two alleles that demonstrated the plasticity of the *AFP* gene family, one encoding 33 AFPs and the other allele only four. In the closely related Pacific halibut, this locus encodes multiple Gig2 (antiviral) proteins, but in the starry flounder, the *Gig2* genes were found at a second locus due to a lineage-specific duplication event. An ancestral *Gig2* gave rise to a 3-kDa “skin” *AFP* isoform, encoding three Ala-rich 11-a.a. repeats, that is expressed in skin and other peripheral tissues. Subsequent gene duplications, followed by internal duplications of the 11 a.a. repeat and the gain of a signal sequence, gave rise to circulating AFP isoforms. One of these, the “hyperactive” 32-kDa Maxi likely underwent a contraction to a shorter 3.3-kDa “liver” isoform. Present day starry flounders found in Pacific Rim coastal waters from California to Alaska show a positive correlation between latitude and *AFP* gene dosage, with the shorter allele being more prevalent at lower latitudes. This study conclusively demonstrates that the flounder *AFP* arose from the *Gig2* gene, so it is evolutionarily unrelated to the three other classes of type I AFPs from non-flounders. Additionally, this gene arose and underwent amplification coincident with the onset of ocean cooling during the Cenozoic ice ages.

## Introduction

Ocean waters freeze near −2 °C, but fish blood and lymph is less salty and freezes at around −0.8 °C (1). Any contact with ice in seawater increases the freezing risk, so some fishes produce antifreeze proteins (AFPs) or antifreeze glycoproteins (AFGPs) (2–4). These AF(G)Ps bind to the surface of ice crystals, preventing growth and decreasing the non-equilibrium freezing point to below −2 °C (5,6). As a result, any internal ice crystals that arise due to contact through the skin, gills or alimentary canal remain small in a quasi-stable supercooled state (7), thereby allowing the fish to live in an icy ecosystem.

Four different types of fish AF(G)Ps, type I, II and III as well as AFGP, occur in species within the clade Percomorpha (Figure 1). Both type I and type III AFPs are restricted to this clade. Type I AFPs are found within four groups within three separate orders (Perciformes (8,9), Labriformes (10) and Pleuronectiformes (11–14)), interspersed with groups producing the three other AFP types. This patchy taxonomic distribution was attributed to convergent evolution of these Ala-rich alpha-helical peptides, but their origins were not known (10,15). Type II AFPs arose from a lectin-like precursor (19), but the presence of this globular, non-repetitive protein in three distantly related fish groups that diverged over 200 million years ago (Ma) (Figure 1), came about via horizontal gene transfer (HGT) (20) rather than convergence. Type III appears to have arisen only once, within infraorder Zoarcales, from a domain of sialic-acid synthase (8–10). Finally, the AFGPs, composed primarily of simple Ala-Ala-Thr repeats where the Thr is glycosylated, arose convergently in northern cods (not shown) and Antarctic notothenioid fishes (such as the Antarctic toothfish) from non-coding DNA and a *trypsinogen* gene, respectively (16–18).

**Figure 1:**
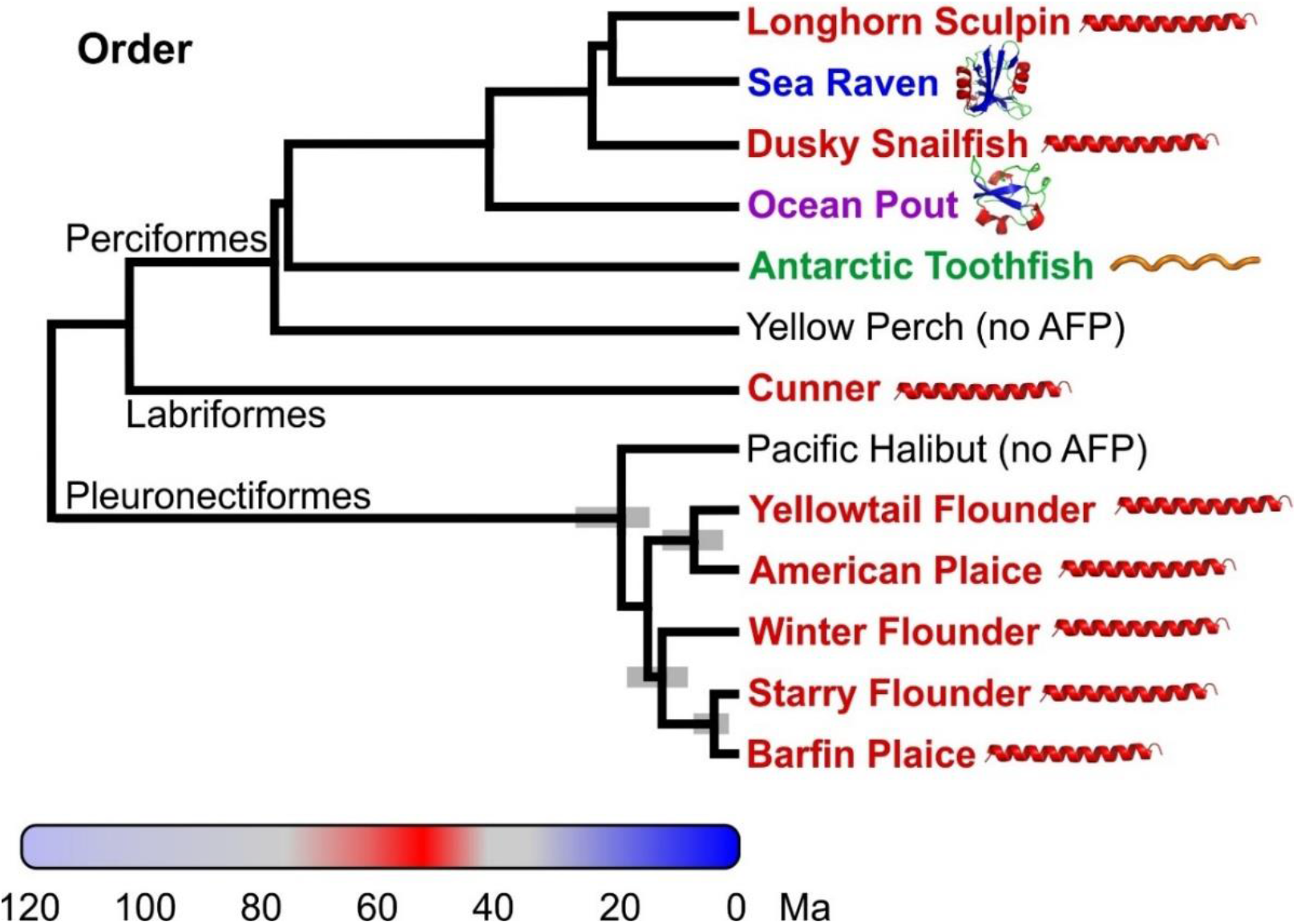
Phylogenetic relationships amongst type I AFP producing fishes and several other species within the clade, Percomorpha, that produce different AFPs (60,61). The common name of species that produce AFPs are coloured red (type I), blue (type II), purple (type II) or green (antifreeze glycoprotein). The 95% highest posterior credibility intervals within the Pleuronectiformes are indicated with grey bars (60). Pacific halibut and yellow perch (black) do not produce AFPs (62). The coloured bar spanning 120 Ma indicates relative ocean temperatures with red corresponding to ice-free oceans and blue corresponding to glacial periods (19). Schematics of the AFP types were generated in PyMOL (63). Binomial names for the species are as follows; *Myoxocephalus octodecemspinosus* (longhorn sculpin), *Hemitripterus americanus* (sea raven), *Liparis gibbus* (dusky snailfish), *Zoarces americanus* (ocean pout), *Dissostichus mawsoni* (Antarctic toothfish), *Perca flavescens* (yellow perch), *Tautogolabrus adspersus* (cunner), *Hippoglossus stenolepis* (Pacific halibut), *Limanda ferruginea* (yellowtail flounder), *Hippoglossoides platessoides* (American plaice), *Pseudopleuronectes americanus* (winter flounder), *Platichthys stellatus* (starry flounder), *Pleuronectes pinnifasciatus* (barfin plaice). Some species were not analyzed in the studies cited above, so the position of the following species with the same genus (dusky snailfish, Antarctic toothfish, Pacific halibut, yellowtail founder, American plaice, barfin plaice) or family (sea raven) was used as a proxy. Other fish, including Atlantic herring and rainbow smelt that are outside Percomorpha, also produce AFPs.

The appearance of these different AF(G)Ps within various groups of fish is correlated with past climate history (Figure 1). After the warming period culminated by the Paleocene-Eocene thermal maximum at 55 Ma (red on color bar), when the oceans were perpetually ice-free (19–21), fish would have had no need of AF(G)Ps for many Ma, and if they were present in prior epochs, they were likely lost. Southern glaciation commenced ~34 Ma (blue on color bar), but continental-scale northern glaciation lagged by ~30 Ma, beginning at ~3 Ma (19). Nevertheless, there is evidence for sea ice and localized ephemeral northern glaciation far earlier, roughly coincident with southern glaciation (19,22). The patchy distribution of AF(G)P types in groups that diverged prior to 20 Ma is consistent with the hypothesis that these proteins arose anew, allowing these species to inhabit a new icy niche as cooling intensified. It is only within recently diverged groups, such as the type I AFP-producing Pleuronectiformes, that AFPs are homologous due to descent from a common ancestor.

Type I AFPs have been best characterized in the winter flounder. There are three isoform classes, all of which are Ala-rich, with Thr appearing at 11 a.a. intervals (15). The abundant small serum isoform HPLC6, produced primarily by the liver (hereafter called a liver isoform), is processed by removal of the secretory signal peptide and pro-region, plus removal of the C-terminal Gly during amidation (23). The mature 37-a.a. peptide is 62% Ala by content and forms a single α helix with three 11 a.a. repeats, delineated by four evenly-spaced Thr residues that lie along one side of the peptide (24). Subsequently, a second class was isolated from skin, hereafter called skin isoforms, although they are expressed in a variety of tissues. These 37-40 a.a. isoforms lack a signal peptide, and their only modification is acetylation of the N-terminal Met residue (25). The third isoform is the much larger hyperactive AFP, hereafter called Maxi, whose only modification is removal of the signal peptide (26,27). This 195 a.a. α-helical peptide folds in half to form an antiparallel homodimer via clathrate water interactions (28).

The presence of type I AFPs in four groups within Percomorpha (Figure 1) could potentially be explained by the presence of the gene in the common ancestor of these groups, followed by its loss in most branches and subsequent gain of different AFPs in a subset of branches. The 76% sequence identity between a winter flounder skin isoform and a longhorn sculpin isoform would seem to support this hypothesis (15). However, other type I AFPs are far less like those from flounders, including the 113-residue dusky snailfish AFP which lacks the 11-a.a. Thr periodicity (8). Additionally, the stark differences in the Ala codon usage in the AFP genes of three of the four groups and the complete lack of similarity of their non-coding sequences led to the hypothesis that they arose via convergent evolution (10,15). Convergence of the AFGPs in northern and southern fishes has been clearly demonstrated following the determination of their progenitors as mentioned above (16–18), but until now such analysis was lacking for any of the type I AFPs.

The starry flounder, *Platichthys stellatus*, is a flatfish that inhabits shallow waters of the Northern Pacific Ocean from South Korea, up through the Bering Sea and down to California, as well as portions of the Arctic Ocean (29,30). It is known to produce type I AFPs, but their sequences were previously unknown (31,32). Loci containing AFP-like sequences were cloned from BAC libraries and both *AFPs* and the progenitor gene, *Gig2* (**g**rass carp reovirus-**i**nduced **g**ene **2**) (33), were identified. Similarity between the loci is restricted to non-coding regions and Gig2 has a different function, related to viral resistance (34). This demonstrates that the AFPs of Pleuronectiformes arose recently and independently of the type I AFPs of other fishes. The two alleles at the *AFP* locus are very different, containing 4 and 33 *AFPs* with Southern blotting demonstrating that gene copy number increases with latitude.

## Materials and Methods

### BAC library construction, screening and sequencing

A BAC (bacterial artificial chromosome) library was constructed by Amplicon Express (Pullman, Washington, USA) from genomic DNA from an individual starry flounder captured off the west coast of British Columbia. All animal studies were conducted in accordance with the Canadian Council on Animal Care Guidelines and Policies with approval from the Animal Care and Use Committee. A total of 12 clones that hybridized to the 3’ untranslated region (UTR) of an AFP transcript were sequenced at the Génome Québec Innovation Centre (Montreal, Quebec, Canada) using the PacBio RS II single molecule real-time (SMRT^®^) sequencing technology (Pacific Biosciences, Menlo Park, California, USA).

### DNA assembly, gene annotation and Southern blotting

The initial assembly was done by the Génome Québec Innovation using the Celera assembler (35). The overlapping regions of different clones were identical except at longer homopolymer or dinucleotide repeat regions. A region containing near-identical 11.2 kb repeats was assembled and evaluated separately, yielding 3.9 assembled repeats out of 12 total, as described in supplementary materials and methods. Genes were annotated using homologs from other fish.

DNA from starry flounders collected at various locations from California to Alaska was Southern blotted and the blots were evaluated using various ^32^P-labelled various probes to *AFP* genes. A more detailed description of all procedures can be found in Supplementary Materials and Methods.

### Nomenclature

Genes are differentiated from proteins using italics. For simplicity, AFPs from starry flounder are named by class with “liver” for small circulating isoforms, “skin” for small isoforms first isolated from skin, “Midi” for an isoform of intermediate size and Maxi for the large circulating isoforms. Numbering is used for classes with multiple isoforms, such as S1 and L1 for the first skin and liver gene at allele 1 respectively. Isoforms from allele 2 are differentiated by letter a (S1a, L1a for example) whereas those from winter flounder are preceded by WF-.

## Results

### Part 1 – Flounder loci

#### Starry flounder AFP genes reside at a single locus

Two BAC libraries made from a single starry flounder caught off Vancouver Island, British Columbia were screened using a probe to the well-conserved 3’ UTRs found in flounder *AFPs*. The tiling paths of 35 positive BACs were determined by PCR screening with a variety of primers (Supplementary Figure 1, Supplementary Table 1) and corresponded to two loci. The first locus was represented by 13 clones and contained five closely spaced *Gig2* genes (Figure 2C) with partial sequence similarity to *AFPs*. Based on the starry flounder genome size obtained from the Animal Genome Size Database (6.5 x 10^8^ bp) (http://www.genomesize.com/index.php) this is consistent with a single gene locus (*Gig2*).

**Figure 2:**
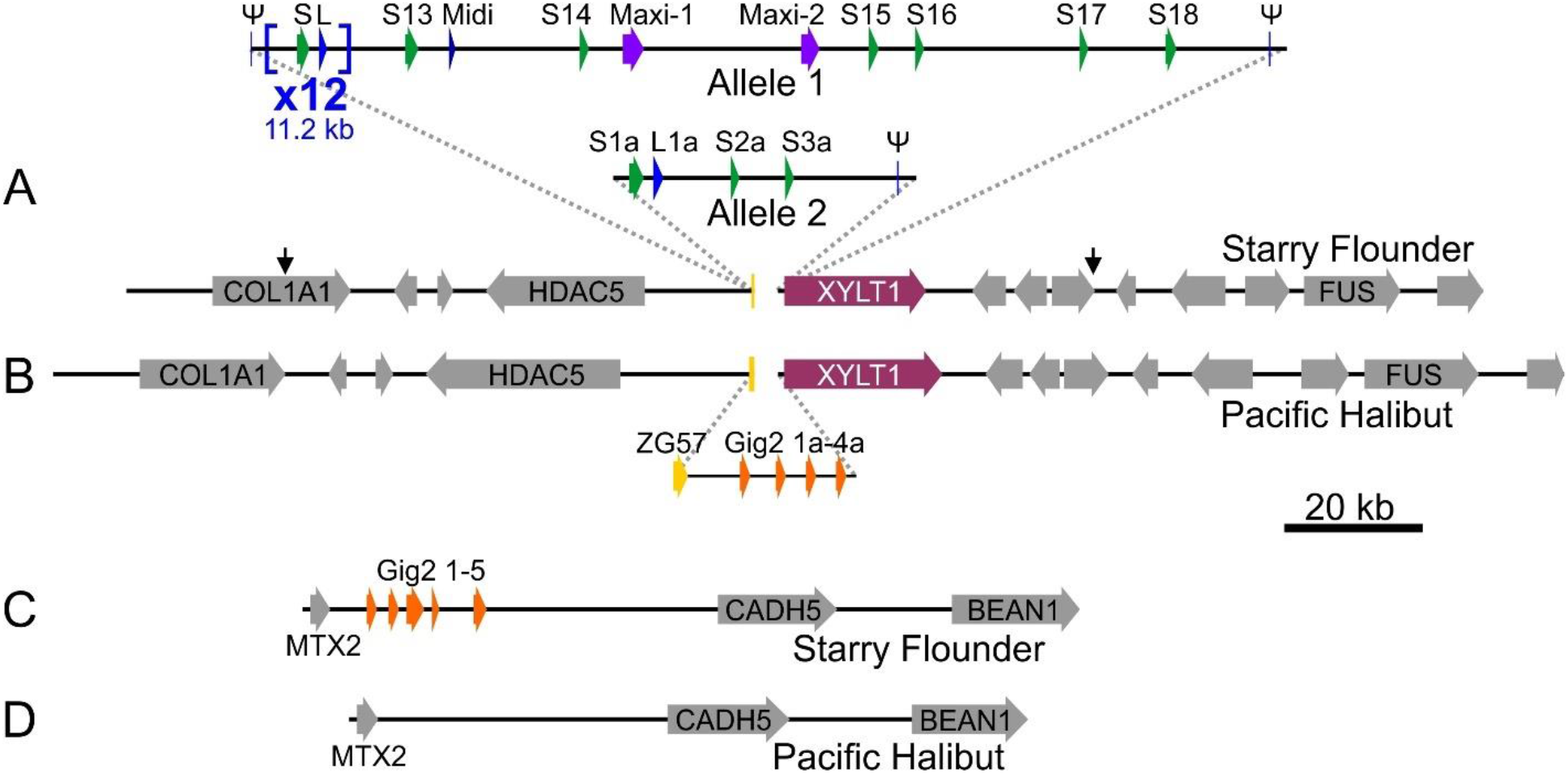
Low-resolution schematic of the *AFP* and *Gig2* loci of starry flounder and Pacific halibut. A solid arrow spans each gene, across all exons and introns, from the start to stop codon, except for *AFP, Gig2* and where non-coding exons were included. Syntenic genes that are not germane to the evolution of the *AFP* are in grey with the acronyms of the shorter genes omitted. All schematics are at the same scale. A) The *AFP* locus from the single fish used to generate the BAC library is shown with the *AFP*-containing segment that differs from Pacific halibut and between the two alleles shown as a pop out. The 33 *AFP* genes in allele 1 are indicated in blue (liver isoforms), green (skin isoforms), dark blue (intermediate length “Midi” isoform) and purple (long “Maxi” isoforms) and are numbered sequentially by type. The first 24 *AFP* genes (12 liver and 12 skin) occur in pairs within twelve nearly identical tandem repeats that are each 11.2 kb in length (shown compressed to one repeat x 12). These are flanked by two short segments (ψ) that are highly similar to portions of the *AFP* genes. The second locus contains four *AFPs* denoted with the suffix “a” and one pseudogene. The black arrows show the boundaries of the locus 2 assembly. B) The segment of Pacific halibut DNA on chromosome 16 that corresponds to the *AFP* locus. The pop out shows the region that differs with respect to the starry flounder locus with the four *Gig2* genes shown in orange. The *ZG57* gene that was partially deleted at this location in starry flounder is in dark yellow and the *XYLT1* gene is in maroon. C) The *Gig2* locus of starry flounder with five *Gig2* genes and D) the corresponding region from chromosome 12 of Pacific halibut. Genes are coloured as above. GenBank accession numbers for these sequences, top to bottom, are OK041463, OK041464, NC_048942 (845791 bp to 1041091 bp), OK041465, NC_048938 (22286642 bp to 22384527 bp).

The remaining 22 clones correspond to two remarkably divergent alleles from a single multigene *AFP* locus (Supplementary Figure 1A & B). This greater number of clones is consistent with the *AFPs* spanning a much larger DNA length (31 or 240 kb) than the *Gig2* locus (17 kb) (Figure 2). The two banks of *AFPs* are allelic as they share the same four flanking genes on each side, including those coding for collagen type 1, α1 (COL1A1) and histone deacetylase 5 (HDAC5) on the upstream side and xylosyltransferase 1 (XYLT1) and RNA-binding protein FUS (FUS) downstream.

#### The two AFP alleles contain a vastly different number of AFP genes

The number of genes within both copies of the locus from this single fish differ greatly as one allele contain 33 *AFPs*, whereas the smaller contains only four (Figure 2A). The difference between the two alleles is not a cloning artefact for two reasons. First, multiple BAC inserts were sequenced for each allele (Supplementary Figure 1A & B), and they were exact matches where they overlapped. Second, the flanking regions of the two alleles are not identical, with around 3% divergence in DNA sequence, primarily within low-complexity regions. However, the protein sequences of the two genes immediately flanking the *AFPs, HDAC5* and *XYLT1* (Figure 2A), are 100% identical.

The structure of the larger allele (allele 1) is complex. Its 33 *AFPs* are flanked on both sides (allele 1) with partial gene sequences (pseudogenes). The single pseudogene in allele 2 is downstream of the four *AFPs* (Figure 2A). Allele 1 contains twelve (Supplementary Figure 2) nearly-identical 11.2 kb tandem repeats, each encoding both a skin and a liver AFP isoform, *L1–L12* and *S1–S12* (Figure 2A, see Nomenclature in Materials and Methods for further details about gene/protein names). These are followed by nine additional *AFPs;* six skin isoforms (*S13-S18*), one longer liver isoform (*Midi*) and two long isoforms (*Maxi-1, Maxi-2*). Allele 2 lacks *Maxi* sequences and contains a single pair of genes encoding a skin and liver isoform (*S1a, L1a*), with high similarity to the pairs within the tandem repeats of allele 1 (Supplementary Figure 3). This region of allele 2 is 94% identical, over 11.9 kb, to the repeat region of allele 1. Two skin isoforms follow, *S2a* and *S3a*, and they closely resemble *S15* and *S16*, respectively. Allele 2 could have arisen from allele 1 via two large deletions, the first removing 11 of 12 repeats through to *Maxi-2*, and the second removing *S17* through *S18*. Alignments between these two alleles can share up to 98% identity over several kb, but all of these contain a few base insertions or deletions in addition to mismatches (not shown). A comparison of the four coding sequences in allele 2 to their closest matches in allele 1 show an average identity of 98.4%.

#### AFP gene structure

All the *AFP* genes, with the exceptions of the pseudogenes that flank the locus, possess two exons (Figure 3, partial data shown), the first of which is non-coding in the case of the *skin* isoforms, but which encodes most of the signal peptide in all other isoforms. The basis for identifying the flanking sequences as pseudogenes are as follows. The 5’ pseudogene of allele 1 lacks a coding sequence but is identical over 80 bp to the 3’-end of the 3’ UTR of the *liver, Maxi* and some *skin* genes. The 3’ pseudogenes of both alleles contain partial coding sequences (16 a.a. or 33 a.a.) that are shorter than the shortest skin isoform (37 a.a.), and the Thr are not spaced at 11 a.a. intervals (Supplementary Figure 4A). Additionally, they lack the first exon due to the insertion of an ~2 kb LINE1 transposon (not shown), which would likely interfere with expression.

**Figure 3:**
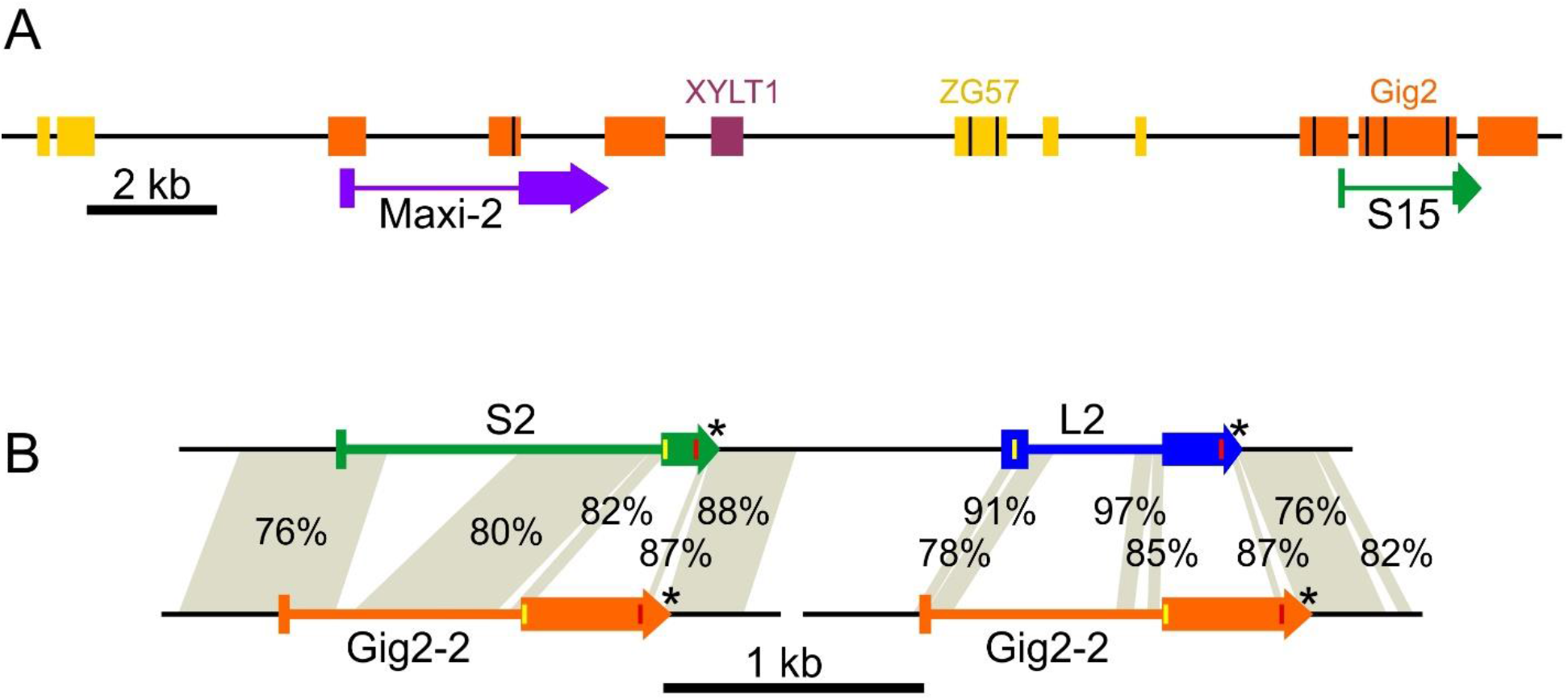
Higher-resolution view of selected *AFP* genes with similarities to *non-AFP* genes indicated. A) A 24 kb segment of Allele 1 containing the *Maxi-2* and *S-13* genes, coloured as in Figure 2, with exons indicated by thicker bars. Blocks show regions of similarity to conspecific *XYLT1* (maroon) and *Gig2* (orange) as well as *ZG57* from Pacific halibut (dark yellow). Black lines within blocks indicate the location of deletions within the *AFP* genes relative to the *non-AFP* genes. Identities range from 70 to 96%. B) Detailed comparison of repeat 2 (bases 87,151–91,650), containing the skin (*S2*) and liver (*L2*) genes, with the *Gig2-2* gene (bases 12,051–14,425). Yellow and red lines within exons represent the start and stop codons respectively and the introns are indicated with a thinner bar. Asterisks denote conserved AATAAA polyadenylation motifs. The shading connects regions of similarity between the two loci with percent identities indicated.

#### There are twelve 11.2 kb AFP-containing repeats in allele 1

The 11.2-kb repeats at the 5’ end of allele 1 were almost identical. By selecting and anchoring the longest reads to polymorphisms in the outer repeats, as described in supplementary materials and methods, the first 2.4 repeats and the last 1.5 repeats were unambiguously assembled. The interior repeats appeared virtually identical, so they were counted using a different method (Supplementary Figure 2). A subset of raw sequence reads, from two clones that overlapped the entire region (BAC45 and BAC182, Supplementary Figure 1) were analyzed. The number of reads corresponding to either the BAC vector or the repeat was compared. The larger BAC45 dataset indicated that there were likely 12 repeats (11.9 ± 0.6), overlapping the estimate of 11 repeats (11.2 ± 0.9) from the smaller BAC182 dataset. The lack of divergence of the internal repeats suggests that they may be undergoing rounds of expansion and contraction through unequal crossing over.

The near identity of the twelve tandem 11.2 kb repeats is mirrored in the protein sequences of the repeats that were assembled. The four liver AFPs (L1, L2, L11, L12) are identical and the last of the three skin isoforms (S12) differs at just one a.a. residue from S1 and S2 (Supplementary Figure 4B & C).

#### The AFPs fall into three main groups

The shortest encoded isoforms are the skin isoforms that lack both a signal peptide and propeptide (Supplementary Figure 4B). Most are 37–39 a.a. long with an acidic residue (Asp) at position 2 and a C-terminal basic residue (Arg) to interact with the helix dipole, as well as three Thr residues at 11 a.a. intervals. The exceptions have a C-terminal extension lacking Arg (S17, S14), a two-residue internal insertion (S14) and both a C-terminal extension and an additional 11 a.a. repeat (S18, 54 a.a.). One winter flounder skin isoform is identical to S3a and a second differs at a single residue (36).

The second group are secreted isoforms that have both a signal peptide and a propeptide that are cleaved from the mature AFP (Supplementary Figure 4C). The starry flounder liver isoforms in the 11.2 kb repeats are 38 residues long after processing, similar in length to the skin isoforms. The liver isoform of the second allele (L1a) has a single Asn mutation at one of the periodic Thr residues. These isoforms have several substitutions relative to their winter flounder counterparts (37,38) and a longer propeptide region. The sequence designated Midi is like the liver isoforms with a signal sequence and propeptide region that are thought to undergo the same N-terminal processing. However, instead of three 11-a.a. repeats, this isoform has six.

The third group are the hyperactive Maxi isoforms (Supplementary Figure 4D), found only in allele 1, where they are adjacent to one another. These isoforms have a signal peptide, but they lack the propeptide domain found in the other liver isoforms. These 194–195 a.a. proteins are over five times longer than most of the skin and liver isoforms and align well with the two known hyperactive isoforms from winter flounder (26,36). The identity between the two starry flounder sequences, Maxi-1 and Maxi-2, is 82%. When compared to the winter flounder sequences, Maxi-1 is more like 5a (82%) than WF-Maxi (79%), whereas the opposite is true for Maxi-2 (79% to 5a vs 84% to WF-Maxi). Maximum-likelihood phylogenetic analysis (Supplementary Figure 5) groups Maxi-1 with WF-5a and Maxi-2 with WF-Maxi, indicating that these two isoforms may have arisen prior to the separation of the winter flounder and starry flounder lineages, over 13 MA ago (Figure 1). This is also consistent with the divergence (18%), between Maxi-1 and Maxi-2.

#### The second cloned locus contains five copies of Gig2

The two BACs that were sequenced (Supplementary Figure 1C) from the *Gig2* locus (Figure 2C) were identical, suggesting they originated from the same allele. The *Gig2* genes lie between the *metaxin-2* (*MTX2*) and *cadherin-5* (*CADH5*) genes, so they reside at a different locus than the *AFP* genes. This locus was isolated because the *Gig2* genes share up to 92% identity to a 252 bp segment of the 3’ UTR *AFP* probe used to screen the library.

The five *Gig2* genes in this locus were identified and annotated by comparison with well-characterized *Gig2* genes from other fishes (33). Gig2 has been shown to protect fish kidney cells in culture from viral infection (34). One of the isoforms (Gig2-4) is 40 residues shorter than the others and may be a pseudogene. The four isoforms that are 147 a.a. long were aligned (Supplementary Figure 6) and they share 73-86% sequence identity. Notably, the sequence of these proteins does not resemble that of the AFPs as they contain little Ala. SMART analysis (http://smart.embl-heidelberg.de/) suggests that residues 20-115 of Gig2-3 are similar to the poly(ADP-ribose) polymerase catalytic domain (expect value of 1.6 x 10^-6^).

### Part 2 – Similar loci in other fishes

#### A syntenic Pacific halibut locus lacks AFPs but contains Gig2 genes

A high-quality genome sequence is available for the Pacific halibut (GenBank Assembly GCA_013339905.1) (39), a species in the same family (Pleuronectidae) as starry flounder. These species shared a common ancestor around 20 MA ago (Figure 1). The region of its genome corresponding to where the *AFP* locus is in the starry flounder shares the same flanking genes on either side, including *COL1A1, HDAC5, XYLT1* and *FUS*, but it completely lacks *AFP* genes (Figure 2B). Instead, it contains four *Gig2* genes. These were annotated in the GenBank deposition (XM_035180664.1) as one combined *Gig2* gene with adjustments for frameshifts. Conspecific transcriptomic sequences in the Sequence Reads Archive database at NCBI (40) were inconsistent with this combined gene model, so they were reannotated to show four copies of *Gig2*, each with a small non-coding exon followed by a coding exon as in the starry flounder *Gig2* genes. The first two genes encode proteins that are highly similar (71-80% identity) to the starry flounder Gig2 proteins (Supplementary Figure 6). The next two contain frameshifts that disrupt the reading frames, so like *Gig2-4* in starry flounder, these may be pseudogenes.

There was one gene found downstream of *HDAC5* in Pacific halibut, just upstream of the *Gig2* genes, that was not found in starry flounder. This gene encodes gastrula zinc finger protein XlCGF57.1 (ZG57), a 56.3-kDa protein that shares no similarity with AFPs. This gene consists of two coding exons and is well conserved in other fishes. However, its location relative to the flanking genes varies. For example, it is found upstream of both *Gig2* and *XYLT1* in yellow perch, but *HDAC5* and the other genes found upstream in flounder and halibut are elsewhere (not shown).

#### The Pacific halibut locus that is syntenic to the Gig2 locus in starry flounder lacks Gig2 genes

The region of the genome in Pacific halibut that corresponds to the *Gig2* locus of starry flounder was also characterized (Figure 2D). Although the flanking genes, *MTX2*, *CADH5* and *BEAN1*, were well conserved, there is a complete absence of *Gig2*-like sequences at this location.

#### Starry flounder AFPs are homologous to AFPs from other Pleuronectiformes

The homology of the winter flounder and starry flounder AFPs is apparent from the similarity of their non-coding sequences. A 2.9 kb portion of a 7.8 kb tandemly-repeated gene from winter flounder encodes a liver isoform (41). Most (88%) of this sequence, which is primarily non-coding, has over 84% identity to the starry flounder 11.2 kb repeat (Supplementary Figure 7). It was not determined if this winter flounder repeat also contained a *skin* isoform.

Additional winter flounder genomic sequences, initially identified as pseudogenes (36), are also highly similar to starry flounder sequences. Two *skin* genes (GenBank accessions M63478.1 (1.4 kb) and M63479.1 (1.2 kb)), are most similar to S14 (Figure 2), with 90% and 85% identity respectively. Additionally, the WF-*5a* gene (GenBank accession AH002489.2) is over 80% identical to both *Maxi-1* and *Maxi-2* over most of its length.

The non-coding sequence of the *AFP* from the more distantly-related yellowtail flounder (Figure 1) (12), is also highly similar to that of the starry flounder *liver* isoform within the repeats. The 5’ UTR (30 bp) is 93% identical and the 5’ UTR is (96 bp) is 96% identical to the *liver* isoforms in the 11.2 kb repeat. Similar comparisons to the non-coding regions of the *type I AFPs* of other orders (Figure 1) failed to identify any similarity (not shown), as was found for comparisons using winter flounder sequences (15).

### Part 3 – The origin of the flounder AFP genes

#### Remnants of three genes indicate that the AFP genes arose at their current location

The region containing the starry flounder *AFPs* was compared to the flanking sequences and to the Pacific halibut *ZG57* locus (Figure 2A & B). A portion of the *ZG57* gene containing the first exon and part of the intron is found just upstream of the first AFP pseudogene in allele 1. This segment encodes 22 a.a. that closely resemble the N-terminal sequence of the halibut protein, but several frameshifts thereafter disrupt the reading frame, and the second exon is absent, so this gene is no longer functional (not shown). Sequences similar to various regions of *ZG57* are found scattered throughout the *AFP* region and some of these are indicated in dark yellow in Figure 3A. Similarly, segments corresponding to the 5’ region of the downstream *XYLT1* gene are also found scattered about, and while only one small segment is found in the region shown in Figure 3A (maroon), three segments totaling 2.2 kb are found within the 11.2 kb repeats (not shown). Some *AFPs*, such as *Maxi-2* (Figure 3A), are flanked by both *ZG57* and *XYLT1* segments. *ZG57* segments are always upstream and *XYLT1* segments are always downstream of *AFPs.* This suggests that a single *AFP* gene arose between *ZG57* and *XYLT1* and that when the *AFP* locus expanded, portions of these flanking genes were duplicated along with the *AFP*.

#### Gig2 was likely the AFP progenitor

A comparison of the *Gig2* and *AFP* loci of starry flounder indicated that there were many stretches of similar sequence, some of which are shown in orange (Figure 3A). As these matches cover a significant portion of the *AFP* gene, except for the coding sequence, this suggests that the *AFP* gene arose from the *Gig2* gene. Furthermore, the greater number of matches to *S15* than to *Maxi-2* suggests that the *skin* gene likely arose first and that subsequent alterations, in which regions similar to *Gig2* were lost, gave rise to the *Maxi* genes.

A more detailed comparison is shown between the skin and liver *AFPs* within the 11.2 kb repeat and the *Gig2-2* locus (Figure 3B). Here again, the skin *AFP* is more like *Gig2* with regions of similarity beginning before and extending across the non-coding exon 1, continuing throughout much of the intron and into exon 2, up to and including the start codon. The coding sequences share no significant similarity, but similarity begins again 31 bp downstream of the stop codon of the *AFPs*, which is right at the stop codon of *Gig2*, and it extends into the 3’ region, overlapping a presumptive poly-adenylation signal. The matches between *Gig2* and the liver *AFP* are more limited, including in the presumptive promoter/enhance region upstream of the gene, and resemble those between *Gig2* and *Maxi-2*.

As mentioned previously, exon 1 of both *Gig2* and skin *AFPs* is non-coding, but for the liver and Maxi *AFPs*, it encodes a signal peptide. This first exon is very well conserved in *S1*, as shown in an alignment to *Gig2-3* (Supplementary Figure 8). It is also reasonably well conserved in *Maxi-2*. A limited number of mutations, such as AGG to ATG to introduce a start codon, along with a small insertion of 23 bases, was sufficient to convert the exon to a signal-peptide encoding sequence. This indicates that the signal peptide arose *in situ*, from the non-coding exon 1 of *Gig2*.

#### Possible origins of the AFP coding sequence

Flounder AFP is Ala rich and these straight α helices provide a flat surface that interacts with ice (24,28). In contrast, Gig2 has a lower-than-average Ala content (~5%), with only one 5 a.a. segment (ACATA, highlighted in cyan, Supplementary Figure 6) that resembles the Ala-rich AFP sequence. For this small segment to give rise to a type I AFP, it should be within a surface-exposed α helix. Fortunately, the structure of a homolog, poly(ADP-ribose) polymerase catalytic domain, is known and the Phyre2 (42) homology model of Gig2 (Figure 4) shows that this ACATA segment is likely surface exposed and is located on the longest helical segment predicted for this globular protein. The AlphaFold2 (43) *de novo* model is very similar and predicts the same surface exposed helix. Deletion of most of the coding sequence, followed by amplification of this short segment, could have given rise to a primordial AFP. Alternatively, a GC-rich sequence encoding numerous Ala residues, such as such as (GCC)_n_, could have replaced the *Gig2* coding sequence.

**Figure 4:**
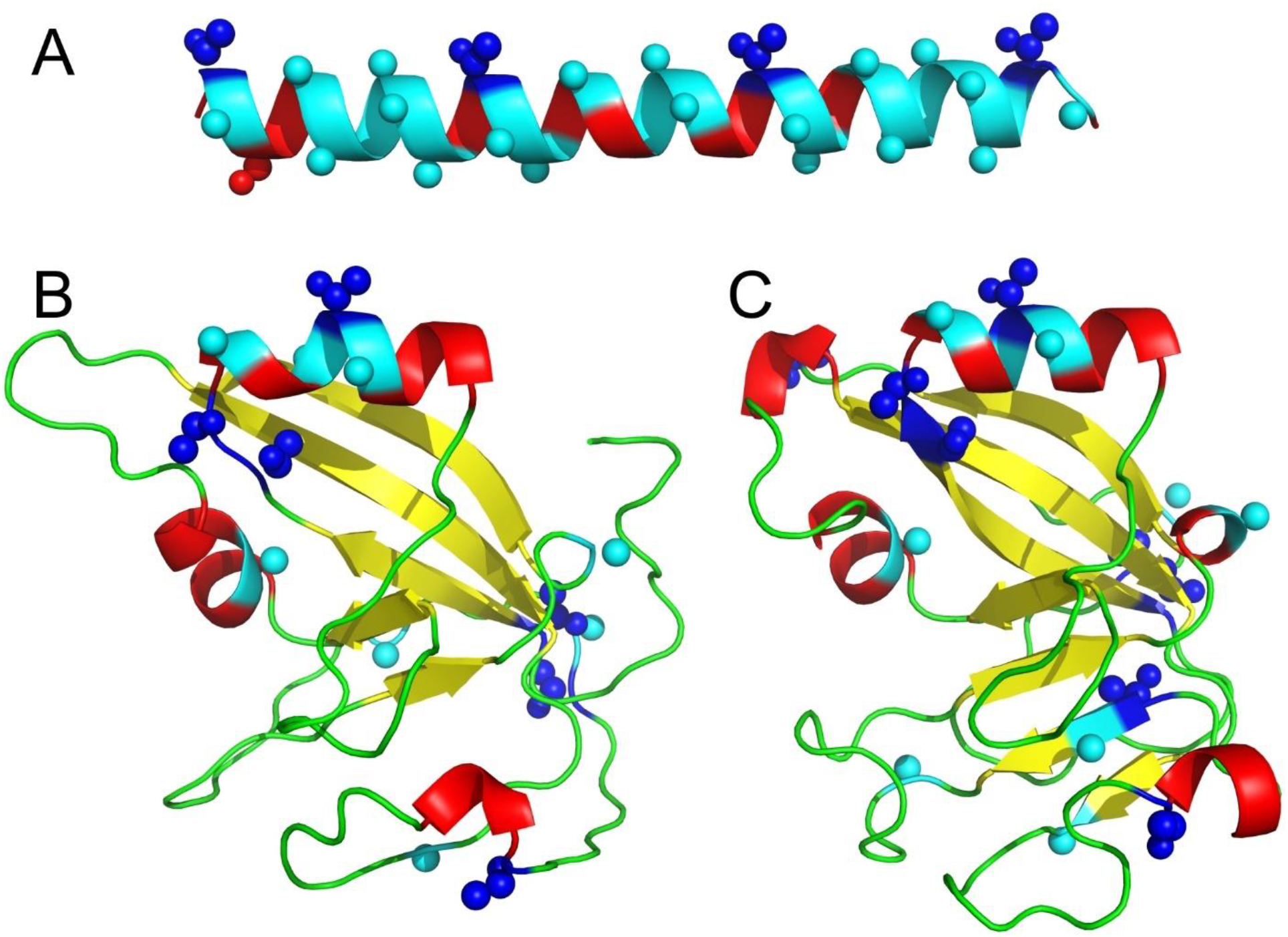
Homology model of Gig2 compared to type I AFP from winter flounder. A) Type I AFP (PDB:1WFA). B) Gig2-3, modelled using Phyre2 (42), was aligned with 100% confidence over 89% of its length to the template PDB:3C4H. C) Gig2-3 modelled without a template using simplified AlphaFold 2.0 (43). The first eight residues (5%) were removed as they were only modelled with low confidence. The images were generated using PyMOL (63) and are shown in cartoon mode with small spheres representing side chains for Ala residues (cyan) and Thr residues (blue). The other residues are coloured by secondary structure with α-helices in red, β-strands in yellow and coils in green.

#### Deduced steps in the generation of the flounder AFPs

The comparisons between the various loci of the starry flounder and Pacific halibut make it clear that the ancestor of the flounder had *Gig2* genes lying between the *ZG57* and *XYLT1* genes (Figure 2A-B). Similar synteny is found in other fishes (not shown). Within the flounder lineage, a gene duplication event led to additional copies of the *Gig2* gene at the second locus, between *MTX2* and *CADH5* (Figure 2C-D). The original *Gig2* genes were then redundant, and one underwent changes that generated a skin *AFP.* This could have come about if the short Ala-containing segment within the α-helix region expanded (Figure 4) or if a segment of repetitive,

GC-rich DNA replaced the coding sequence. The gene was then duplicated an unknown number of times, and the non-coding exon 1 evolved to encode a signal peptide (Supplementary Figure 8). Further gene duplications and gene losses (Supplementary Figure 3), as well as expansions and contractions of the repetitive coding sequences, gave rise to the extant complex alleles due to the selective pressure (or lack thereof) of living around sea ice.

#### Allele 2 is more prevalent in starry flounders from warmer waters

The fish that was used to construct the library, and which had the two differing *AFP* alleles, was caught in southerly Canadian waters of the North Pacific, off the western side of Vancouver Island (pink/green circle, Figure 5A). In contrast, a genomic Southern blot of four fish collected from the Haida Gwaii, approximately 300 km further north (location 1, Figure 5A), showed that the larger *AFP* allele 1 was prevalent at this location (Figure 5B-2). Two intense bands, corresponding to the *skin* and *liver* genes within the 11.2 kb repeat, confirm the repetitive nature of this repeat. Bands corresponding to the predicted sizes of all the other genes from allele 1 were also observed, further confirming the accuracy of our assembly. A more detailed analysis of the correspondence between these bands and the two *AFP* alleles is shown in Supplementary Figure 10. There is some evidence of limited polymorphism as a few unexplained bands were present in one or two of the fish, but all these fish appear to be homozygous for alleles very similar to allele 1, as bands corresponding to the unique and well-separated fragment sizes expected for *S2a, S3a* and *S4a* were not observed.

**Figure 5:**
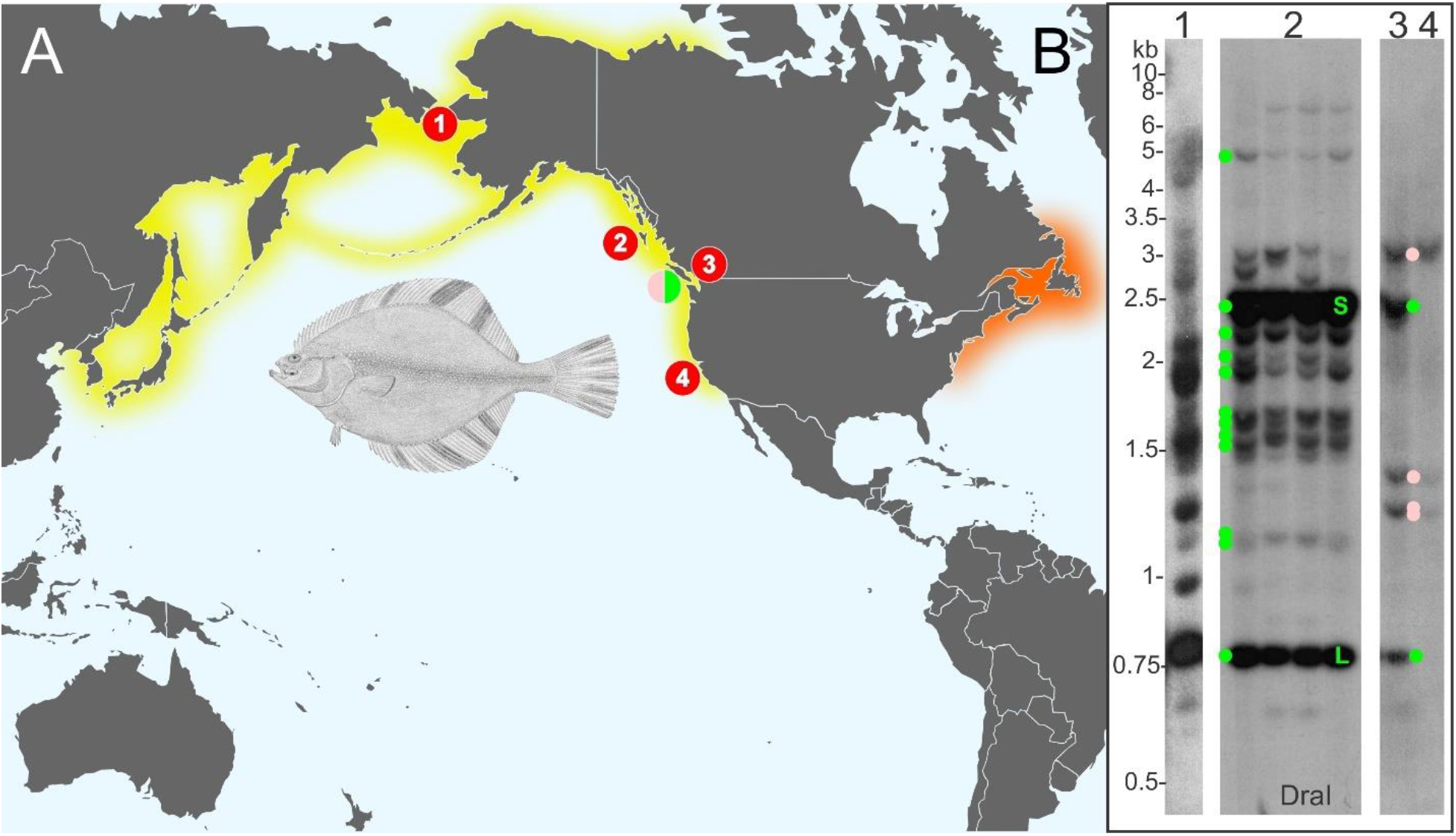
Southern blot of genomic DNA from starry flounder collected from various regions throughout its range. A) The native range of starry flounder and winter flounder are indicated with yellow and orange shading respectively. The locations where fish were harvested for Southern blotting are indicated with the red numbered circles while the location of the fish harvested off Vancouver Island used to generate the BAC library is indicated with the split pink/green circle. B) Southern blot of DNA digested with *DraI* for individual fish from the Bering Straight, Alaska (1) English Bay, British Columbia (3), Monterey Bay, California (4) and from four fish from Haida Gwaii (2). The blots were probed with a sequence from allele 1 (bases 77,759 to 77972) that overlaps the second exon of the skin *AFP* within the 11.2 kb repeat. The expected locations of fragments from allele one are indicated by green dots with S (skin) and L (liver) for the genes within the repeats. Pink dots correspond to fragment sizes expected to arise from allele 2. The starry flounder drawing is a public domain image by H. L. Todd and the map was modified from an image by Dmthoth, CC BY-SA 3.0, both of which were obtained from Wikimedia Commons.

In contrast to the large *AFP* copy number of the more northerly starry flounder, a fish caught in Monterey Bay, California (location 4, Figure 5B-4), only has bands consistent with allele 2. Although at a similar latitude as the sequenced flounder from the west coast of Vancouver Island, the fish caught in the warmer slightly brackish waters of English Bay, off Vancouver (location 3, Figure 5B-3), had bands consistent with allele 2, along with some moderately intense bands consistent with the *skin* and *liver* genes within the 11.2 kb repeats. We speculate that it contains an allele similar to allele 2 that still has a small number of 11.2 kb repeats remaining. A fish from Alaska (Figure 5B-1), approximately 1500 km further north from Haida Gwaii, had many intense bands with sizes that were not consistent with either allele. Together, these results suggest that gene copy number is correlated with risk of ice exposure and that numerous alleles with differing numbers of *AFP* genes can be found within this species.

## Discussion

Taxonomically restricted genes (TRGs) confer phenotypic novelty on their hosts and the selective pressures of new environments often provide the driving force for their development (44,45). For example, water striders have colonized the water surface due in part to TRGs that generate a “fan” on the middle leg that provides propulsion across the surface (46). Similarly, the climate cooling that intensified during the latter half of the Cenozoic Era generated an icy sea environment that had been absent for at least tens of Ma (19,22), and which would have excluded fish from shallow water niches where ice is found until the *AFP* genes arose in certain species, including the recent ancestors of the starry flounder. These and other TRGs arise in a variety of ways (44), including via duplication and divergence of existing genes, as for example with AFGP, type II and type III AFP (17,47,48), or *de novo* from non-coding DNA (AFGP (16,18)). It can be difficult to determine the mechanism, as selection for a new function can lead to rapid divergence, erasing the similarity to the progenitor sequence (49). This erasure likely occurred with the coding sequence of the flounder *AFP* gene as it bears little similarity to the *Gig2* progenitor. Fortunately, the *AFP* arose recently, so extensive similarity between the flanking regions of the two genes was retained (Figure 3). Additionally, the lineage-specific duplication of the *Gig2* genes at a second locus, as well as sequential duplications of segments of the flanking genes at the original locus (Figures 2 & 3), shows that the *AFP* gene arose, *in situ*, at the original *Gig2* locus via gene duplication and divergence.

It is now clear that the AFPs of Pleuronectiformes, such as starry flounder, are not homologous to the type I AFPs found in the other three lineages (snailfish, cunner and sculpin) within Perciformes and Labriformes, as these other *AFPs* lack similarity to *Gig2*. It was proposed that the snailfish AFP could have arisen from a frameshifting of the Gly-rich region of either keratin or chorion cDNAs that were inadvertently cloned along with the *AFP* genes (50). However, the similarity did not extend into non-coding segments. As all these genes arose within the last ~20 Ma, they would be expected, like the flounder’s, to retain some evidence of their origins in their non-coding regions, since diversifying selection would be lower here. Currently, the origin of the three other type I AFPs remains unknown.

The convergence of the AFPs from four lineages to Ala-rich helices, sometimes with Thr residues at 11 a.a. residues (9,10,15,25), suggests that this motif is well-suited to interacting with ice. Similar convergence, albeit with a different structural framework, was seen with arthropod AFPs that adopt a beta-helical conformation. A beetle (yellow mealworm) and a fly (midge) produce tight, disulfide-stabilized solenoids, with an ice-binding surface composed of a double row of Thr residues or a single row of Tyr residues, respectively (51,52). The looser solenoid of the moth (spruce budworm) is more triangular and lacks bisecting disulfide bonds, but like the beetle AFP, its ice-binding surface consists of a double row of Thr residues (53). This suggests that there are nascent structures with propensities to evolve into AFPs, but that different types are more likely to arise in marine versus terrestrial environments because of the vastly different requirements for freezing point depression.

When a novel gene arises from a pre-existing one, non-coding sequences are thought to be almost as important as coding sequences (54). It is likely that the promoter and enhancer sequences controlling expression of the *Gig2* gene were co-opted, for two reasons. First, the *skin* genes and *Gig2* share high identity upstream of the first exon. Second, the expression patterns of *Gig2* in zebrafish (33) and the winter flounder skin *AFPs* (25) are similar as they are expressed in a variety of tissues. The tissue- and season-specific enhancement of the liver *AFPs* (55) may have arisen later, given that its gene lacks similarity to the upstream regions of the *Gig2* gene. However, all the genes retain the two exons and the polyadenylation signal.

The rapid divergence of the starry flounder *AFP* coding sequence from the *Gig2* progenitor is reminiscent of that observed for the *AFGP* that was derived from the *trypsinogen* gene (17). For the *AFP*, a 15 bp segment, corresponding to 5 a.a. in a helical region of the protein, was likely retained and amplified (Figure 4). For *AFGP*, the amplified segment was only 9 bp long and it overlapped the acceptor splice junction at the start of exon 2. Both gene types retained the first exon, which is non-coding in skin *AFPs* and *Gig2*, but which encodes a signal peptide in both *AFGP* and *trypsinogen*. However, the first exon of the flounder *liver, Midi* and *Maxi* genes does encode a signal peptide and similarity with the *Gig2* non-coding exon shows that it arose, *in situ*. This is reminiscent of the origin of the signal peptide of type III AFP (47), where an additional 54 bp in exon 1 gained coding potential, generating a signal peptide.

The number of *AFP* genes was higher in starry flounders from the northern waters of Alaska and British Columbia than in flounders from more southerly waters (Figure 5). Variation in gene copy number was also observed in winter flounder from different regions along the Atlantic coast, with animals from warmer waters having fewer genes (56). The same pattern has been observed for ocean pout, which can have up to ~150 genes that produce type III AFP (57). As many of the *AFP* genes are arranged in tandem arrays, they are likely prone to rapid expansion and contraction via unequal crossing over (58), providing variation that would be subject to environmental selection.

Gene duplication also provides additional copies that can undergo neofunctionalization (58), which is how the three main classes of type I AFPs found in flounders (Maxi, liver and skin) arose. The properties of these isoforms differ dramatically as Maxi is far more active than either the skin or liver isoforms (27), and expression of the liver isoform is extremely high in this tissue (59). Unequal crossing over likely led to the loss of the *Maxi* genes and the majority of the *skin* and *liver* genes in the shorter starry flounder AFP allele. A similar process may have occurred in the American plaice. Despite being closely related to the yellowtail flounder that possesses both liver and Maxi isoforms (12,14,60) (Figure 1), American plaice serum only contains Maxi-like AFPs (14). This suggests that the common ancestor of both of these fish had the liver isoform and that the plaice locus may have undergone contraction, losing the small liver-specific *AFP* genes. Similar processes, working on a smaller scale, may also be responsible for the generation of isoform variation. For example, liver-like isoforms with extra copies of the 11-a.a. repeat are found in both starry flounder (Midi with three extra repeats) and yellowtail (one extra repeat (12)). This plasticity may also explain why the banding pattern from the Alaskan starry flounder observed by Southern blotting is so different from that of fish from Haida Gwaii (Figure 5), despite both having large numbers of *AFP* genes.

In summary, the origin of the flounder *AFP* from the gene encoding the globular, antiviral Gig2 protein, via gene duplication and divergence, has been determined. Detailed comparisons between the two loci elucidate the steps involved in the evolution of the *AFP*. Although the flounder AFP is superficially similar to the type I AFPs of other groups, all of which are extended alanine-rich alpha-helical proteins of varying length, it clearly arose by convergent evolution. The two extended loci that were characterized from starry flounder encode either the *AFP* genes or five of the *Gig2* progenitor genes. The two *AFP* alleles sequenced contain either four or 33 *AFP* genes, indicating that gene copy number can vary dramatically. These genes encode skin, liver and Maxi *AFPs*, with the number of *AFP* genes being higher in fish that inhabit colder waters.

## Acknowledgements

We thank Eric Clelland, Dave Riddell, and other staff at the Bamfield Marine Sciences Centre, Bamfield, BC for collecting and shipping starry flounder blood. Amplicon Express (Pullman, WA) made two BAC libraries and corresponding nylon filters. The McGill University and Génome Québec Innovation Centre provided high-quality PacBio sequencing and assembly services without which this project would not have been possible. We are grateful to Nick Ostan for mapping the BACs and to Gary K. Scott and Kyra K. Nabeta for providing genomic DNA and Virginia K. Walker for comments on the manuscript. This work was funded by CIHR Foundation award FRN 148422 to PLD, who holds the Canada Research Chair in Protein Engineering.

## Abbreviations

(AFP): Antifreeze protein
(AFGP): antifreeze glycoprotein
(AF(G)Ps): antifreeze proteins and antifreeze glycoproteins
(BAC): bacterial artificial chromosome
(UTR): untranslated region
(Gig2): grass carp reovirus induced gene 2
α1 (COL1A1): collagen type 1
(HDAC5): histone deacetylase 5
(XYLT1): xylosyltransferase 1
(FUS): RNA-binding protein FUS
(Ma): million years ago or megaannum
(MTX2): metaxin-2
(CADH5): cadherin-5
(ZG57): gastrula zinc finger protein XlCGF57.1
(TRGs): taxonomically restricted genes
(HGT): horizontal gene transfer

