## Supplemental Information for "Origin of the type I antifreeze gene in flounders in response to Cenozoic climate change"

**Supplemental Information for "Origins of the type I antifreeze gene in flounders in response to Cenozoic climate change"**

**Laurie A. Graham, Sherry Y. Gauthier and Peter L. Davies**

### Supplementary Materials and Methods

*BAC library construction, screening and DNA purification*

Adult starry flounder were captured off the west coast of Vancouver Island (49°N, 125°W) by staff and students of the Bamfield Marine Sciences Centre, British Columbia under the supervision of Eric Clelland. Blood was collected from the caudal vein of an individual fish and collected in 4.0 mL lavender BD Vacutainer® tubes containing $K_2EDTA$ (BD, Mississauga, Ontario, Canada) and flash frozen. The library was constructed by Amplicon Express (Pullman, Washington, USA). Briefly, DNA was extracted and partially digested with either *HindIII* or *BamHI* prior to ligation into the pCC1BAC vector (Epicentre, Madison, Wisconsin, USA) transformed into DH10b T1 phage-resistant *E. coli* (Life Technologies, Grand Island, New York, USA). Over 66,000 clones with average insert sizes of 90 to 105 kb respectively were obtained. Nylon filters with clones arrayed in duplicate were screened using a $^{32}$P-labelled probe amplified from genomic DNA, encompassing bases 122348 to 122723 of AFP locus 1 GenBank accession OK041463) which corresponds to the well-conserved 3′ untranslated region (3′ UTR) found in all the flounder AFP genes. Additionally, it is non-repetitive and has a balanced GC content of 48%, whereas the repetitive coding sequences are >70% GC. A total of 35 positive clones were obtained and categorized based upon the fragment sizes generated using various primers specific to the 3′ UTR of the AFP gene and seven were selected for sequencing. The remaining clones were screened by PCR using additional primer pairs based upon these clones and another five clones were selected for sequencing. BAC DNA was prepared using the Qiagen Large-Construct Kit (Qiagen, Toronto, Ontario, Canada) as per the manufacturer's instructions but with the addition of a proteinase K digest (100 µg/mL at 50 °C for 1 h) following endonuclease digestion as well as phenol/chloroform extractions and dialysis following the final step.

*DNA sequencing and assembly*

Sequencing and initial assembly was done by the McGill University and Génome Québec Innovation Centre (Montreal, Quebec, Canada) using the PacBio RS II single molecule real-time (SMRT®) sequencing technology (Pacific Biosciences, Menlo Park, California, USA). Long-read (10 or 20 kb) libraries were generated and read numbers varied from 60 million to over 500 million per BAC clone. The assembly was performed by the Celera assembler (1). The long PacBio reads were corrected using HGAP (Hierarchical Genome Assembly Process). The assemblies generated from overlapping clones were identical barring length variations at longer homopolymer and dinucleotide repeat regions. For these, a series of N's were inserted to approximate the length uncertainty.

The only region which failed to assemble reliably contains nearly identical 11.2 kb repeats. The ends of this region were verified and extended by combining long sequence reads (> 15 kb) from overlapping clones. These reads were screened by BLAST (2) for BAC vector ends and/or for unique indels (insertions or deletions) unique to the terminal repeats, as well as for quality throughout their length. They were trimmed to the unique sequence segments to ensure they assembled in the correct register using the contig editor from GeneStudio Professional Edition V 2.2.0.0 (http://genestudio.com/). The assembly was manually edited to the point where only four long reads remained. To ensure accuracy from such low coverage, the repeats were assembled to each other and to the CCS (circular consensus sequence) reads to ensure that all polymorphisms were represented in >5% of the higher-accuracy CCS reads. Polymorphisms were not considered in isolation but were verified by their co-occurrence within individual CCS reads.

The number of 11.2 kb repeats was counted using the sequence datasets for BAC45 and BAC182 as these two clones spanned the repeat region (Supplementary Figure 1). The Galaxy (https://usegalaxy.org/) FASTA/FASTQ tool “Filter sequences by length” was used to select reads that were between 1–2 kb in length (3) and these were divided into bins of 1000 sequences each. There were seven 1-kb query sequences from the BAC vector selected to exclude repetitive sequence, and seven 1-kb query sequences from the 11.2 kb repeat selected to match only one region within the repeat. The number of hits (≥ 75% identity over 300 bp) for each BLASTn comparison of query to binned sequence were counted. Values were divided by the average number of hits to the BAC vector to normalize the BAC vector hits to unity and to obtain an estimate of repeat number. The average and standard error of the mean for the seven values (correcting for those repeat sequences present in the 3′ region of either BAC45 or BAC182 one additional time) of both the BAC vector and repeat were determined and the SEM was calculated using the quotient rule for errors.

*Gene Annotation*

Genes were initially identified by BLASTn and BLASTx (https://blast.ncbi.nlm.nih.gov/Blast.cgi) using overlapping 10 kb segments of each construct that were masked for simple repeats using Repeat Masker (http://www.repeatmasker.org/) (4) and transposable elements using Censor (http://www.girinst.org/censor/index.php) (5). The gene structure was determined by comparison to the corresponding loci from other fish using Genewise (http://www.ebi.ac.uk/Tools/psa/genewise/) (6) or tBLASTn (2) and annotated using SnapGene (https://www.snapgene.com/). The assembled and annotated loci were deposited under GenBank accession numbers OK041463-OK041465.

*Analysis of the Pacific halibut genome*

BLAST searches using starry flounder sequences were performed on the halibut genome (GeneBank Assembly GCA_013339905.1). GenBank flatfiles encompassing the genomic regions that are syntenic with those from starry flounder were downloaded and visualized using SnapGene (https://www.snapgene.com/). One group of genes (*Gig2*) were misannotated, so more detailed comparisons were done based on homologous sequences from the protein, EST and a conspecific SRA database (liver transcriptome, GenBank accession SRR11826188) identified through BLAST searches (2).

*Southern blotting*

Starry flounder were collected by trawling from several locations across their range. The southernmost fish was collected in Monterey Bay, California (37°N, 122°W) and tissue (liver) was provided by Gary Scott, while the northernmost were collected from Alaska, in the Bering Strait (65°N, 168°W). Another five were collected in British Columbia; one from English Bay, Vancouver (49°N, 123°W) and four from Haida Gwaii (52°N, 131°W).

Genomic DNA was extracted from tissue (liver or muscle) as described in Scott *et. al*. (7)The DNA was digested with either *AseI* or *DraI* (NEB) and approximately 5 µg of genomic DNA was loaded per lane on a 1% agarose gel. The DNA was transferred to Zeta-Probe GT membrane and hybridized to $^{32}P$-labelled skin (*S1*) probe (bases 77,759 to 77972) or liver (*L1*) probe (79,680 to 79,983) of GenBank OK041463, which includes 21 bp of upstream sequence, the entire coding region and 79 bp of 3′ untranslated region) according to manufacturer's instructions. The final wash was done at 65°C, 0.1XSSC, 0.1% SDS for 25 min.

**Supplementary Results**

*BAC libraries and PacBio sequencing was instrumental in assembling these large, complex loci*

A genomic library of Starry flounder DNA was previously constructed in Lambda DASH II and two clones were sequenced using Roche 454 technology (data not shown). Each clone contained only two widely spaced AFP genes and there were zero reads spanning the GC-rich coding sequence of the longer AFP gene. Whole genome sequencing was impractical as highly repetitive gene loci generally fail to assemble properly. Therefore, a large insert BAC library was generated, and selected BACs were sequenced using PacBio SMRT (single molecule real time) sequencing (~85% accuracy at the time) with lengths up to ~20 kb. Higher accuracy circular consensus sequences (CCS) were generated from shorter fragments sequenced multiple times. The lack of an PCR steps, as well as the characteristics of the polymerase, ensure that both repetitive and GC-rich regions were sequenced. One caveat is that the number of bases within longer homopolymeric or dinucleotide repeats was not accurately determined.

Hindsight revealed that this approach was optimal, given current technology, for several reasons. First, each locus/allele contains multigene families with 4 to 33 members and long stretches of near identical sequence. For the *Gig2* locus, the longest segment is 2.4 kb with 94% identity and in the shorter *AFP* allele, there are 1.4 kb segments with 97% identity as well as 1.7 kb segments with 91% identity. The exceptionally long sequencing reads obtained from the PacBio instrument were able to span these repeats, unlike traditional Sanger sequencing or most next-generation sequencing technologies. Second, the *AFP* coding sequences are very GC rich. PacBio had no difficulty sequencing though the ~0.6 kb coding sequences of the Maxi isoforms that are 77% GC, unlike the Roche 454 technology. Third, the use of BACs enabled us to ascertain that the single fish used to build the library contained two *AFP* alleles with very different gene copy numbers. This would likely have confounded any whole genome assembly. The one quandary that was the region containing the twelve 11.2 bp repeats was not completely assembled.

*Confirmation of the structure and allelic nature of the two banks of AFP genes*

The repetitive nature of the *AFPs* within the two alleles could potentially lead to recombinational cloning artefacts, but as all but a very small region of allele 1 (center of BAC135) was sequenced across two distinct BACs, this is very unlikely (Supplementary Figure 1). The sequence of overlapping regions of the BAC clones matched exactly, except where there was uncertainty in the number of nucleotides within homopolymeric or dinucleotide repeat regions. In addition, the regions of high similarity between all *AFPs*, excluding those within the 11.2 kb repeats, were short enough to be spanned by PacBio reads (Supplementary Figure 9). The many small indels (insertions and deletions) within these matches cannot be resolved at the scale of this dot plot, but cumulatively, they cause the apparent angle of some matches to deviate from 45°.

Further confirmation of the allelic nature of these loci was provided by the flanking regions (Figure 2A). These sequences were identical between clones corresponding to the same allele, barring uncertainty in stretches of mono- or di-nucleotide repeats. Yet the overall identity between alleles was only 97%, with ~ 2/3$^{rds}$ of the differences within indels, predominantly within low-complexity repeats. These genes were annotated with between three and 49 exons (not shown), whereas the *AFPs* have only two (Figure 3). The coding sequences themselves are less divergent as, for example, the protein sequences of the two HDAC5 alleles are identical.

*The 11.2 kb repeats were counted by BLAST and CCS reads confirm their near identity*

The high similarity between the entirety of the 11.2 kb repeats was far greater than the similarities between the downstream *AFP* genes and for this reason, they could not be assembled in their entirety. However, the number of repeats was counted by comparing the number of reads that matched various portions of the vector to the number matching the repeat, giving estimates of 11.9 ± 0.6 for BAC45 and 11.2 ± 0.9 for BAC182 (Supplementary Figure 2). The near identity of the internal repeats was confirmed by aligning ~100 high-quality CCS reads to one repeat and scoring the variations. As there are twelve repeats, sequencing errors present in one or two reads could be immediately discounted. Those that remained were accounted for in repeats 1, 2 and 10. The coding sequences, other than the one difference mentioned for S10, were 100% identical.

*Flounders from Haida Gwaii possess AFP alleles very similar to allele 1.*

A detailed comparison of DNA of the four flounders from Haida Gwaii, digested with two different restriction enzymes, showed bands consistent with the predicted sizes of all 33 *AFPs* from allele 1 (Supplementary Figure 10). The bands corresponding to the genes within the 11.2 kb repeat (*L1-12* and *S1-12*) are intense, as expected, and although they hybridize to both probes, their intensity is greater with the matching probe. Those genes with bands that overlap the liver bands in the AseI digest (*S13*, *S14* and *Midi*), are visible as separate bands in the DraI digest. The unlabelled bands, most of which differ between fishes, could arise due to gain or loss of restriction sites, gene duplications or insertions and deletions. This indicates that there is allelic variation, even among fish caught at the same location.

**Supplementary Table 1**: PCR primers used to determine the extent of the various BAC clones as indicated in Supplementary Figure 1.

| Specificity | Name | Sequence 5' to 3' |
|---|---|---|
| One site in allele 1 and 2<br><br>Named for allele 1 | BAC45_5'_F | GTGCAGATAAACCAGGTAGGATGA |
| | BAC45_5'_R | GTGACTGACCAGGAGTAGTTAACA |
| | BAC102_5'_F | CGAAGATCACTCCCAAGTTCC |
| | BAC102_3'_F | CAGTGCAGCATTCGACACAAT |
| | BAC102_3'_F | TCAGGACCAGAATAGCTTTGCAT |
| | BAC102_3'_R | GACTGACACTGGCTTCAAACTTC |
| Specific for allele 1 | BAC45_R1_F | AGTGATGGAGCGGAGACAAGGAGAT |
| | BAC45_R1_R | GATCACTTCGTCACTGCTTCTCCTA |
| | BAC45_3'_F | TGGATGTCTGTACATGTGTTTGG |
| | BAC45_3'_R | TTAACAACACGTTGGAACACCTG |
| | BAC132_5'_F | CCAAGTAGAATCTCCTCCTTCCG |
| | BAC132_5'_R | AGATTGGAGGTGATTGTCATGGC |
| | BAC135_Mid1_F | TGCTAGGCACGGTCTAATACAG |
| | BAC135_Mid1_R | GAGTGGTAAAGGCGATCACAA |
| | BAC135_Mid2_F | GGGCTTTGCTTTATTCCAGTTG |
| | BAC135_Mid2_R | GGGTTAAGTGCTGGCTTCATAG |
| | BAC135_3'_F | CTTGACTAGCCAGAACCACCGTT |
| | BAC135_3'_R | GGCAGGCGTTTATCGCATCTA |
| Specific for allele 2 | BAC79_F | CACATTGAATCTGTCGTTACCTCG |
| | BAC79_R | GGTGGTTGAGCTCGTTCTTAAAGT |
| Multiple sites in Allele 1, one site in allele 2 | BAC45_GEN_REPa_F | CACGAGGGGAACGAAGGTTA |
| | BAC45_GEN_REPa_R | TCATGGGAACCGGGATTTTGGTC |
| | BAC45_GEN_REPb_F | CGTGTCCATGACCTAGTTTCAGAAG |
| | BAC45_GEN REPb_R | TAACGTTCACTCTGACTCACAAGA |
| Gig2 | BAC79,122_5'_F | TGGAGCTGGTCAACAACATGCT |
| | BAC79,122_5'_R | TCAGAGGCCACGAGTACGGAC |
| | BAC79,122_3'_F | CAGCCTGTTGATACAAATCCGTCTC |
| | BAC79,122_3'_R | AGTCTCCCTGTCCAGTGTGTT |

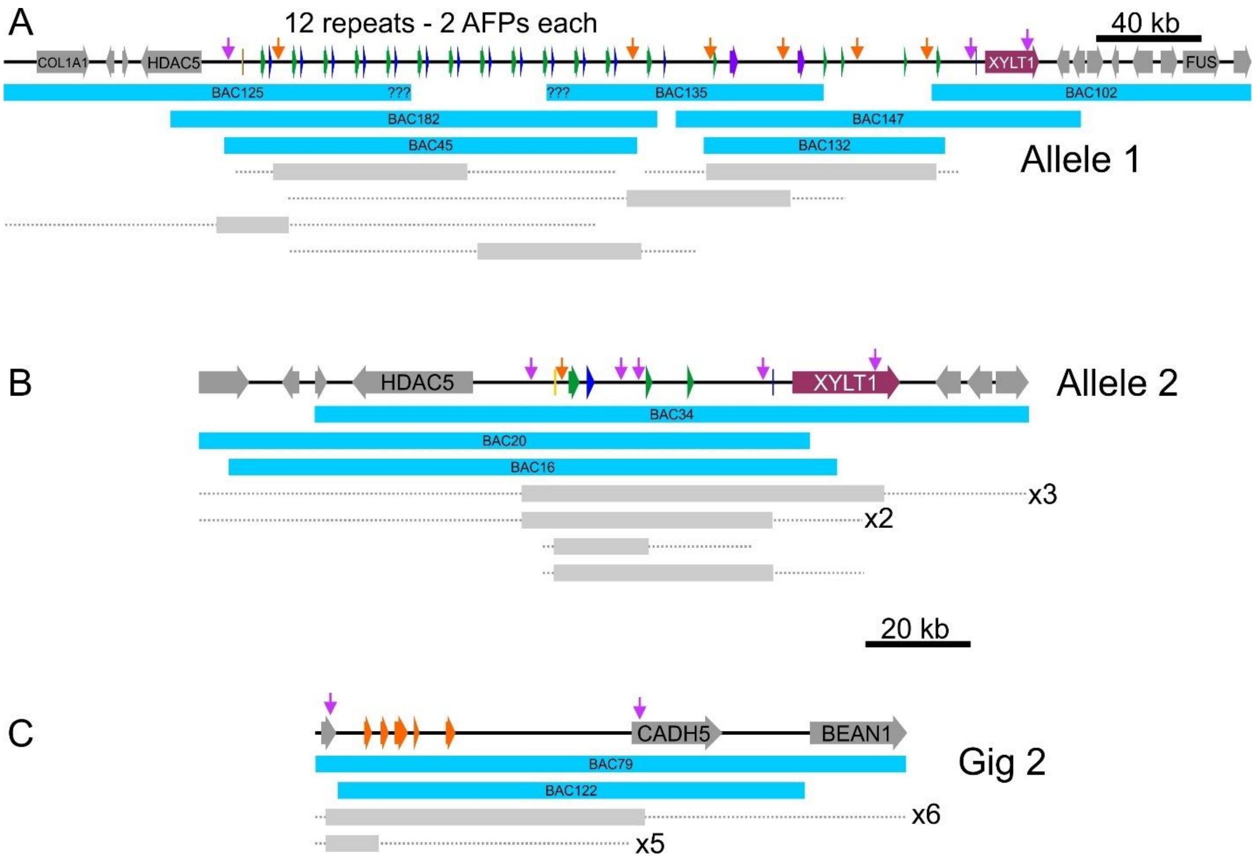

**Supplementary Figure 1:** Schematic diagram of BAC clones which overlap A) AFP allele 1, B) AFP allele 2 and C) the Gig2 locus. Genes are coloured as in Figure 2 and the deduced number of tandem repeats is indicated for allele 1. The sequenced BAC clones are indicated with cyan bars. The span of other BAC clones (grey bars with dashed lines indicating uncertainty) were determined by PCR using location-specific primers (purple arrows) and primers that were location and allele specific (orange arrows) (Supplementary Table 1). All clones were PCR positive using primers specific to the 3′ UTR found in both the Gig2 and AFP genes.

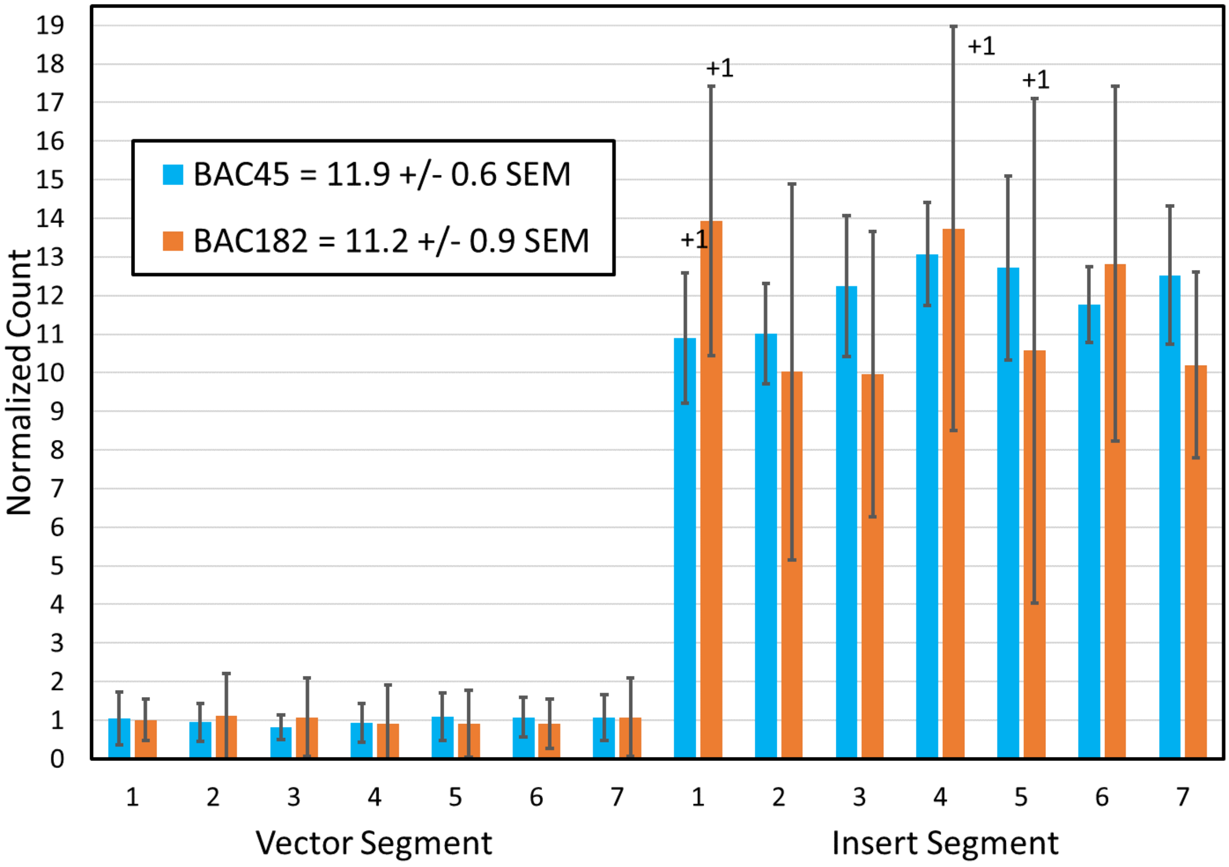

**Supplementary Figure 2:** Determination of the number of 11.2 kb repeats in two clones (BAC45 in blue, and BAC182 in orange) that span the entire repeat region, (Supplementary Figure 1). Raw sequence reads that were 1-2 kb long were divided into eight (BAC182) or twelve (BAC45) groups of 1000. The number of BLAST hits (≥ 75% identity over at least 300 bp) within each group, to 1 kb stretches corresponding to seven non-repetitive regions of the single-copy BAC vector and seven unique regions within the repeat, were counted. Values were normalized to copy number by dividing all counts by the average number of hits to the BAC. The error bars indicate standard deviation. Some segments are also found once, downstream of the repeats, and these are indicated with +1. The estimate of the number of repeats and the standard error of the mean (SEM) are shown. The BAC45 dataset was larger and of better quality, with ~4 times the number of hits, so it provides a more accurate estimate of repeat number.

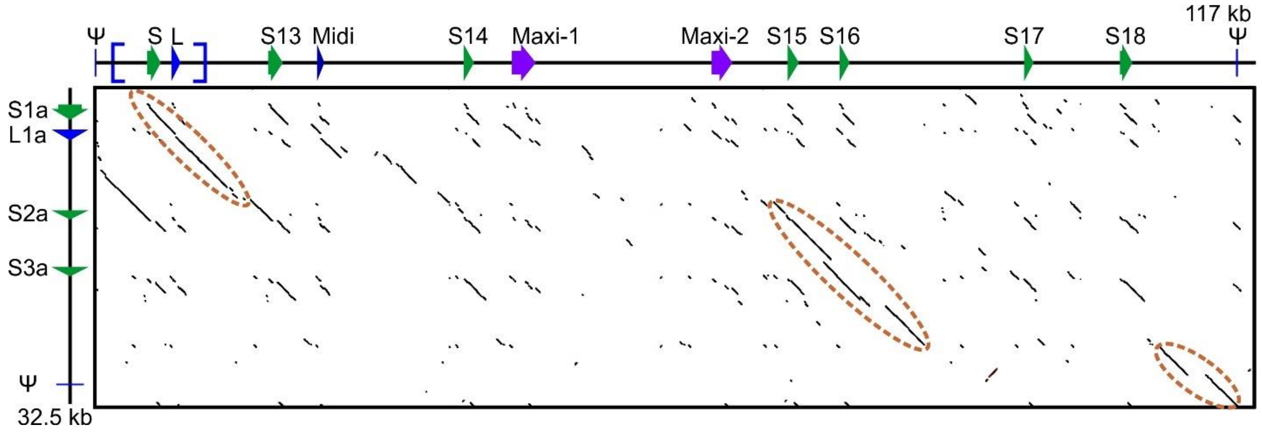

**Supplementary Figure 3:** Dot plot comparison of the *AFP*-containing regions of allele 1 (horizonal axis) and allele 2 (vertical axis) with forward matches in black and reverse matches in red, generated using YASS (8). Regions of highest similarity are indicated with brown dashed ellipses and the gene schematics are from Figure 2

## A

```
Ψ2-Allele 1     MDAATIAAAAA-----------------TATAT 16
Ψ-Allele 2      MDAATIAAAAAAATAAKAAEDAASATAAAATAT 33
                ***********                 :****
```

## B

```
S1,2,1a         MDAPARAAAATAAAAKA--AAEAT-----------KAAAAKAAAATKAAR-------- 37
S13             MDAPAKAAAATAAAAKA--AAEAT-----------KAAAAKAAAATKAAR-------- 37
S12             MDAPARAAAATAAAAKA--AAEAT-----------AAAAAKAAAATKAAR-------- 37
S17             MDAPARAAAATAAAAKA--AAEAT-----------AAAAAKAAADTKAAAAAAAAL-- 43
WF-S1           MDAPAKAAAATAAAAKA--AAEAT-----------AAAAAKAAAATKAGAAR------ 39
S16             MDAPAKAAAATAAAAKA--AAEAT-----------AAAAAKAAAATKAGR-------- 37
S3a,WF-S2       MDAPAKAAAATAAAAKA--AAEAT-----------AAAAAKAAAATKAAAAR------ 39
S14             MDAPAAAAAATAAAAKAKAAAEAT-----------AAMAAKAAAATKAAAALLGSWAS 47
S2a             MDAPAAAAAATAKAAKA--AAEAT-----------AAAAAKAAAATKAAAAR------ 39
S15             MDAPAAAAAATAAAAKA--AAEAT-----------AAAAAKAAAATKAAAAR------ 39
S18             MDAPAAAAAATAAAAKA--AAEATAAAAAAAAAATAEAAAKAAAATKAAAAAAAAR-- 54
                ***** ****** ****  *****                ****** ***.
```

## C

```
                                          ↓                         ↓
L1,2,11,12      malslftvgqliflfwtirineaNPDPAAKAAAVADPAAAAVAPAADAFSAAADTASDAAAAAAA
L1a             malslftvgqliflfwtirineaNPDPAAKAAAVADPAAAAVAPAADAFSAAADTASDAAAAAAA
WF-L1           malslftvgqliflfwtmriteaSPDPAAKAA----PAAAAA-PAA----AAPDTASDAAAAAAL
WF-L2           malslftvgqliflfwtmriteaNPDPAAKAV-----------PAA----AAPDTASDAAAAAAA
Midi            malslftvgqliflfwtiriteaNPDPAAKAAAAA-----------DAFSAAADTASDAAAAAAA
                ***************** ** ** *******                     ** ***********

L1,2,11,12      T---------------------------------AAAAKAAAEKTARDAAAAAAATA----RG 91
L1a             T---------------------------------AAAAKAAAEKNARDAAAAAAATA----RG 91
WF-L1           T---------------------------------AANAKAAAELTAANAAAAAAATA----RG 82
WF-L2           TAATAAAAAAAT----------------------AATAAKAAALTAANAAAAAAATAAAAARG 91
Midi            TAATAAAAAAATAATAAKAAAATAATAKAAAAATAATAAAAAMATIEAAAKAAAATAAAA-RG 116
                *                                 ** *  **      ** ******    **
```

## D

```
Maxi-2          malslftvgqiiflfstiriteaNIDPAAKAAAAAAAAAAAATAADAAAAAATIAASAAA 60
WF-Maxi         malslftvgqfiflfwtisiteaNIDPAARAAAAAAASKAAVTAADAAAAAATIAASAAS 60
Maxi-1          mthslftvgqfiflfwtisitea-IDPAAQAAAAAAAAAAVVTAADAAAAAANAAANAAA 59
WF-5a           malslftvgqfiflfwtisitea-IDPAAKAAAAAAAATAVVTAAAAAAAAAAIAATAAA 59
                *: *******:**** ** **** *****:*******: *..*** ******  **.**:

Maxi-2          VAAATAADAAAASIATINAAAAAATATAAAAAIAAKETATAAASAAAAASAAAAAAQATI 120
WF-Maxi         VAAATAADDAAASIATINAASAAAKSIAAAAAMAAKDTAAAAASAAAAAVASAAKALETI 120
Maxi-1          VAAATAADVATASIATIKANAAAAAATAAAAAIAAEEAATAAATAAAAAAATAATAQAAI 119
WF-5a           VAGATAADAAAASIASINANTAAAAAIAAAAAKAAEEAAATAAAAAATTAATAATAQATI 119
                **.***** *:****:*:* :*** : ***** **:::*::**:***:: *:** *   :*

Maxi-2          NDKAAAAAATTATTAAAAAAATATTAAAAAAATATIENAAATKAAAAASAVAAAAVTAAT 180
WF-Maxi         NVKAAYAAATTANTAAAAAAATATTAAAAAAAKATIDNAAAAKAAAVATAVSDAAATAAT 180
Maxi-1          FDKAAAAASTTATTAATAAAATATTAAAAAAATETIDKAAAAAAAAAATAVATAAAAAAT 179
WF-5a           KDKAAAAAASTATNAAAAAAATATTAAAAAVAKTTIDKAAAAVAVAAATAVAAAAATAAT 179
                  *** **::**..**:*************.*. **::***: *.*.*:**: **.:***

Maxi-2          AAATAAATLEAAAAKAAAVAVSAAAAAAAAAIAAAAA-- 217
WF-Maxi         AAAVAAATLEAAAAKAAATAVSAAAAAAAAAIAFAAAP- 218
Maxi-1          AAATAAATLGAAAAKAAATAVAAAAAAAIAAAAAAAAPP 218
WF-5a           AAATAAATLGAATVKAAATAVNAAAAAAAATAAAAAAPP 218
                ***.***** **:.****.** ****** *: * ***
```

**Supplementary Figure 4:** Alignments of the AFP sequences of starry flounder along with selected sequences from winter flounder. Variable a.a. are highlighted yellow (variation 1) or grey (variation 2) where they occur. A) The translations of the remnant coding sequences of the two presumptive pseudogenes at the 3′ end of the AFP alleles. B) Skin isoforms, including two from winter flounder (WFs1, WFs2, GenBank accessions M63479.1 and M63478.1 respectively). C) Liver isoforms, including two from winter flounder (WFL1, WFL2, GenBank accessions M62416 and DQ062445 respectively) with the secretory signal peptide in lowercase font and the pro-peptide region in italics. Arrows indicate cleavage sites. Underlining indicates the residue interrupted by the intron (phase 2). D) Hyperactive isoforms, including two from winter flounder (WF-Maxi, WF-5a GenBank accessions EU188795.1 and AH002489.2 respectively) with the signal peptide and intron location indicated as above.

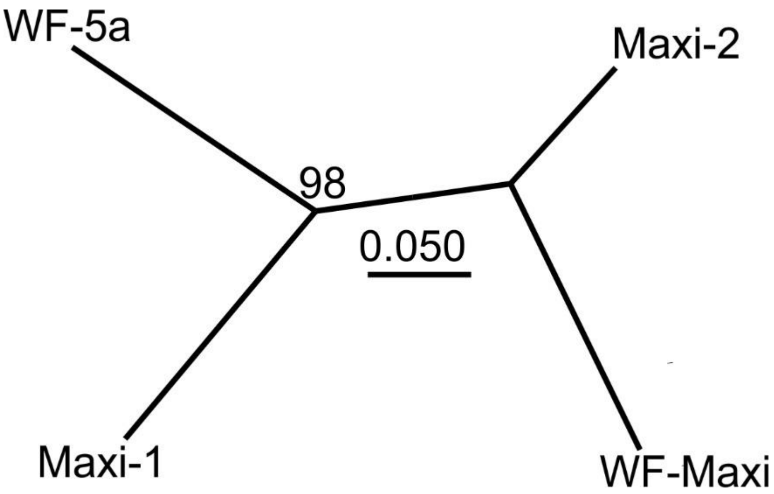

**Supplementary Figure 5:** Unrooted maximum-likelihood phylogenetic tree generated from the alignment in Supplementary Figure 3D using MEGA V10.1.8 (9). The Whelan and Goldman plus frequencies model with six gamma categories was selected using the model test and 100 bootstrap replicates were performed. The scale bar represents 5% divergence and the bootstrap value (100 replicates) is indicated at the node.

```
SF-Gig2-1       -----MYWAEDDFDPPPGTFRLGLAQPVDYQTYVMYHGTTRECATSILNSGFRQSLDGML 55
PH-Gig2-1a      MTCTNMKWAEDDFDPPPGTFRLGPSQPVSDQTYVMYHGTTSQNSRAIQNSGFRRSSGGML 60
PH-Gig2-2a      MTCTNMKWAEDDFDPPPGTFRLGPSQPVSDRTYVMYHGTTSQNSRAIQNSGFRRSSGGML 60
SF-Gig2-3       -----MYWAEDDFDPPAGTFRLGLSQPVNQRTYVMYHGTTRACATAIRKSGFRQSSGGML 55
SF-Gig2-2       -----MKWAEDDFVLPPGSSRLGLAQPVNDLIYVMYHGTTRACATDIQKSGFRPSSDGML 55
SF-Gig2-5       -----MKWAEDDFDLPPGTYRLDLSQPVDGRTYVMYHGTTRACATAIQKSRFRKSSDGML 55
                     * ******  * *: **. :***.   ********   :  * :* ** * .***

SF-Gig2-1       GCGVYLSRDLEKASRYPIDCPEYDRIIIKVLVQVGKVICIDHQNHQFQKIWPYYGYDTAH 115
PH-Gig2-1a      GPGVYLSRDLQRASRYPIDHPEHDKVVIKVKVRVGKVIAINHQYHPSQKTWQYQGFDTAW 120
PH-Gig2-2a      GPGVYLSRELRKASRYPIDHPEHDRVVIKVKVRVGKVIAINYQNHQFQTTWHDQGFDTAW 120
SF-Gig2-3       GRGVYLSRDLEKACRYPLDHPEYDRVVIKVKVRVGKVIAINRQNHPFRETWYDHGYDTAW 115
SF-Gig2-2       GCGVYLSRDLQKASRYPINHPEYDRVVITVLVHVGKVIAINRQKHPLQKTWHDHGYDTAW 115
SF-Gig2-5       GRGVYLSRDLQKASRYPIDHPEYDRVVIKVKVRVGKVIAINRQLHPLQKTWHDHGYDTAW 115
                * ******:*.:*.***:: **:*:::*.* *:*****.*: * *  :  *   *:***

SF-Gig2-1       VPPHCGMVPSGLEEDCVWDPRRIQIIQIFNAT   147
PH-Gig2-1a      VPPNCGMVKSGLEEDCVWDPARIQILQIIKPT   152
PH-Gig2-2a      VPPNCGMVKSGLEEDCVWDPARIQILQIIKPT   152
SF-Gig2-3       VPADCGMVKSGLEEDCVWDPARIKIIQIIKPT   147
SF-Gig2-2       VPPGCGMVPSGLEEDCVWDPTRIQILKIINPT   147
SF-Gig2-5       VPPGCGMVPSGLEENCVWDPARIEILKIINPT   147
                **  **** *****:*****  **:*::*:: *
```

**Supplementary Figure 6:** Characteristics of the Gig2 progenitor protein. A) Alignment of Gig2 isoforms from starry flounder (SF) and Pacific halibut (PH). The segment within a predicted α helix (Figure 4) that is most similar to the AFPs is highlighted in cyan. Isoforms SF-Gig2-4, PH-Gig2-3a and PH-Gig2-4a are not included as their coding sequences are disrupted by one or more base deletions. Variable a.a. are highlighted yellow (variation 1) or grey (variation 2 to 3) where they occur and Ala residues are in red.

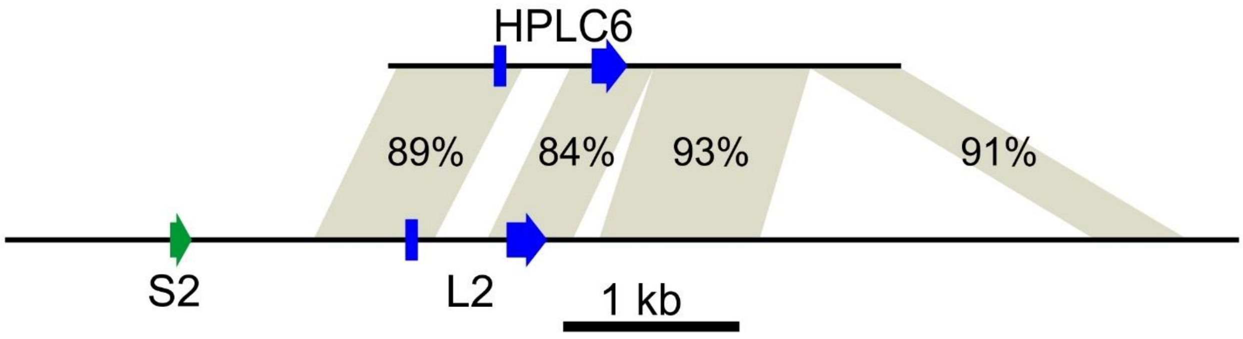

**Supplementary Figure 7:** Similarity between the longest winter flounder *AFP* clone (*HPLC6*, GenBank accession M62415.2) and a segment of the second 11.2 kb repeat of starry flounder containing a skin (S2) and liver (L2) gene. Thicker segments represent the coding sequences. The shading connects regions of similarity between the two loci with percent identities indicated.

A

```
Gig2-3     ACTCGAAGTGAACGTTTATAAGGGCCACTCCCTAAAAGTTTTCAGAAGGACTCACACACT 60
           ||| ||||||||  |  ||||| ||  |||||||||||||||||  ||||||||||||||
S1         ACTTGAAGTGAA--TAAATAAGAGCTGCTCCCTAAAAGTTTTCATCAGGACTCACACACT 58

Gig2-3     GTTCACTGTCGCACACTAAGGTACGTGAACACTCACTTTGTTTCTCCTACAAATCTGGTT 120
            |||||||||| ||||| |||||||||||||||||||||||||||||||||||||||| |
S1         TTTCACTGTCGAACACTCAGGTACGTGAACACTCACTTTGTTTCTCCTACAAATCTGG-T 117

Gig2-3     TTACTGTAAATATCTCGGGAAGGAGTGAAGGATATCTGCATTATCCCCGAGGGGC  175
           ||||||||||||||| ||||||||  ||||||||||||||||||||| |||||||
S1         TTACTGTAAATATCTTGGGAAGGAAGGAAGGATATCTGCATTATCCCAGAGGGGC  172
```

B

```
                                                          R  T  H  T  L
Gig2-3     ACTCGAAGTGAACGTTTATAAGGGCCACTCCCTAAAAGTTTTCAGAAGGACTCACACACT  60
           ||| || |||||| ||||||||| |  |||||| | |||| ||| || | ||| | ||||
Maxi-2     ACTTGACGTGAACTTTTATAAGGTCTGCTCCCTTA-AGTTCTCAAAATGGCTCTCTCACT  59
                                                          M  A  L  S  L
             F  T  V  A  H                        *
Gig2-3     GTTCACTGTCGCACAC----------------------TAAGGTACGTGAACACTCACTT  98
            |||||||||| |||                       | ||||||||||||||||||||
Maxi-2     CTTCACTGTCGGACAAATCATTTTCTTATTTTCGACAATCAGGTACGTGAACACTCACTT  119
             F  T  V  G  Q  I  I  F  L  F  S  T  I  R

Gig2-3     TGTTTCTCCTACAAATCTGGTTTTACTGTAAATATCTCGGGAAGGAGTGAAGGATATCTG  158
           ||||||| ||| |||||||||||| |||||||||||| ||||||||  ||||||||||||
Maxi-2     TGTTTCTTCTATAAATCTGGTTTTGCTGTAAATATCTTGGGAAGGAAGGAAGGATATCTG  179

Gig2-3     CATTATCCCCGAGGGGC  175
           ||||||||| |||||||
Maxi-2     CATTATCCCAGAGGGGC  196
```

**Supplementary Figure 8:** Alignment between the exon 1 containing regions of *AFPs* and *Gig2-3*. Exonic sequences are highlighted yellow. A) Alignment to *S1*, these segments are 90% identical and the exon shown is non-coding. B) Alignment to *Maxi-2* with the sequence of the signal peptide shown, along with a translation of the corresponding region of the non-coding Gig2 exon. These segments are 76% identical.

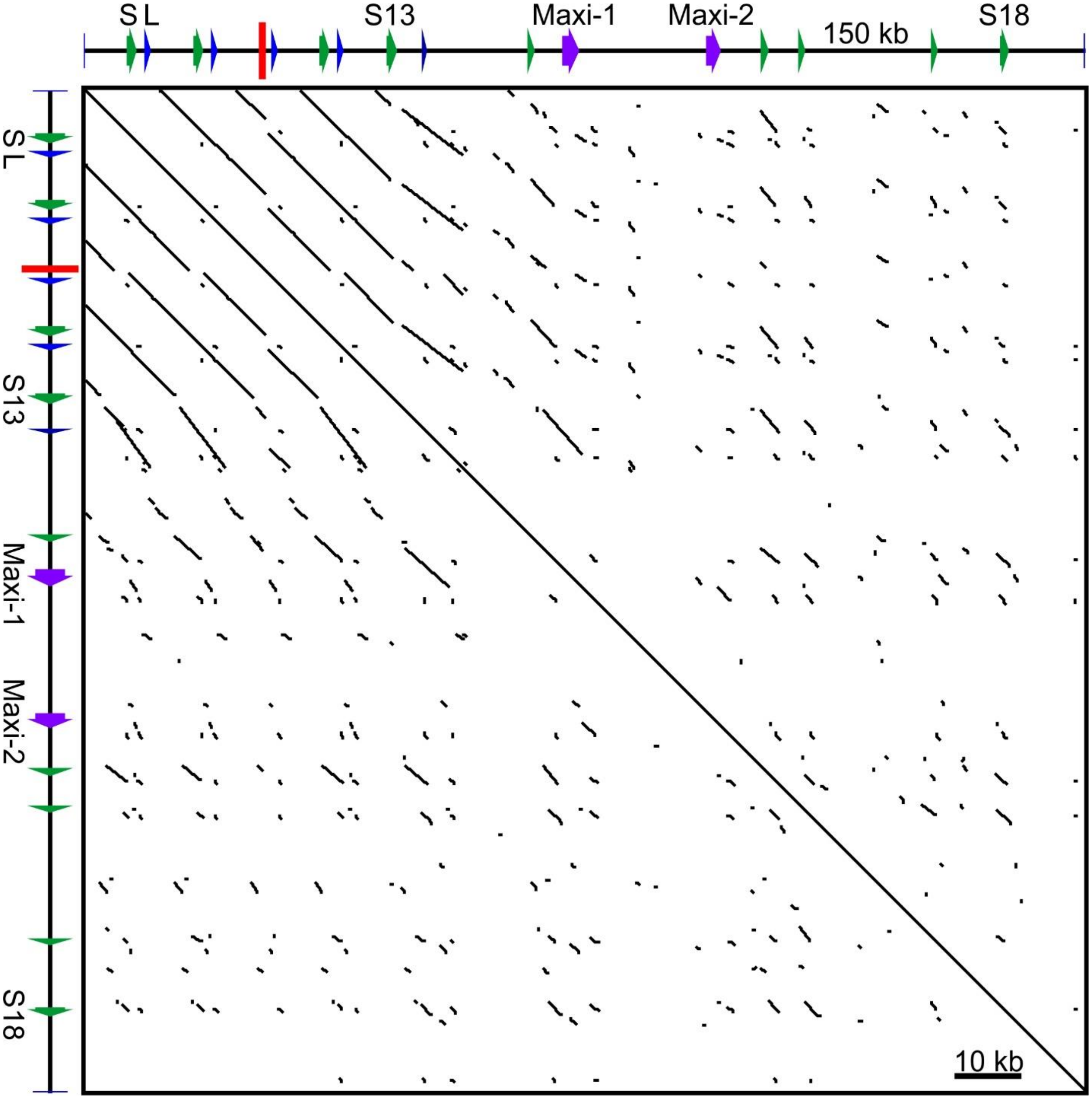

**Supplementary Figure 9:** Dot plot comparison of the *AFP*-containing region of allele 1 compared to itself generated using YASS (8). The gene schematic is from Figure 2 but here all sequenced portions of the 11.2 kb repeats are included. The red rectangle indicates the gap containing 8.1 repeats, which is flanked by 2.4 repeats and 1.5 repeats on either side. The repeats are 99% identical over their 11.2 kb lengths and their longest match outside the repeats is to a 3.8 kb segment overlapping S13, with 97% identity.

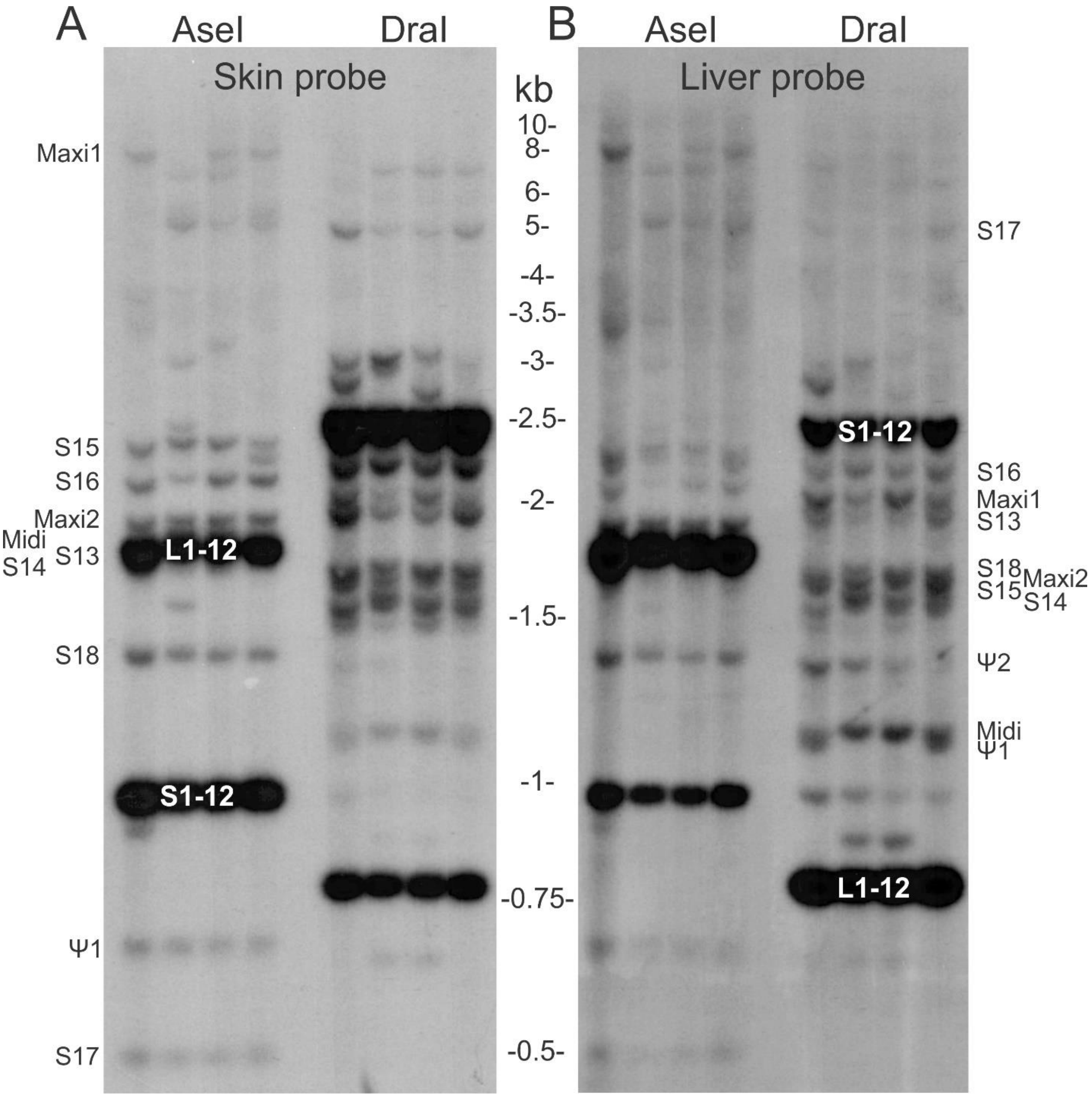

**Supplementary Figure 10:** Southern blot of genomic DNA from four starry flounders collected near Haida Gwaii, digested with *AseI* or *DraI*. A) Blot probed with a fragment from skin-1, with the predicted locations for a theoretical *AseI* digest of allele 1 labelled on the left. B) The blot was stripped and reprobed with a fragment of liver-1 with the labels for the theoretical *DraI* fragments on the right.
